# Scuphr: A probabilistic framework for cell lineage tree reconstruction

**DOI:** 10.1101/357442

**Authors:** Hazal Koptagel, Seong-Hwan Jun, Joanna Hård, Jens Lagergren

## Abstract

Cell lineage tree reconstruction methods are developed for various tasks, such as investigating the development, differentiation, and cancer progression. Single-cell sequencing technologies enable more thorough analysis with higher resolution. We present Scuphr, a distance-based cell lineage tree reconstruction method using bulk and single-cell DNA sequencing data from healthy tissues. Common challenges of single-cell DNA sequencing, such as allelic dropouts and amplification errors, are included in Scuphr. Scuphr computes the distance between cell pairs and reconstructs the lineage tree using the neighbor-joining algorithm. With its embarrassingly parallel design, Scuphr can do faster analysis than the state-of-the-art methods while obtaining better accuracy. The method’s robustness is investigated using various synthetic datasets and a biological dataset of 18 cells.

**Author summary:** Cell lineage tree reconstruction carries a significant potential for studies of development and medicine. The lineage tree reconstruction task is especially challenging for cells taken from healthy tissue due to the scarcity of mutations. In addition, the single-cell whole-genome sequencing technology introduces artifacts such as amplification errors, allelic dropouts, and sequencing errors. We propose Scuphr, a probabilistic framework to reconstruct cell lineage trees. We designed Scuphr for single-cell DNA sequencing data; it accounts for technological artifacts in its graphical model and uses germline heterozygous sites to improve its accuracy. Scuphr is embarrassingly parallel; the speed of the computational analysis is inversely proportional to the number of available computational nodes. We demonstrated that Scuphr is fast, robust, and more accurate than the state-of-the-art method with the synthetic data experiments. Moreover, in the biological data experiment, we showed Scuphr successfully identifies different clones and further obtains more support on closely related cells within clones.

## Introduction

Reconstructing cell lineage trees, from single-cell data, for healthy tissue is a fundamental computational problem with enormous potential for studies of development and differentiation [1–6]. There are two related reconstruction problems for cancer tumors: reconstruction of clonal trees and the reconstruction of tumor phylogenies from single cells. Several data types, e.g., bulk DNA, single-cell DNA, and single-cell RNA, have been used for the latter two problems [7–11]. All these reconstruction methods exploit mutations and attempt to reconstruct trees in which the proximity between a pair of cells, or clones, is correlated with the similarity between their patterns of mutations. The somatic mutation rate in humans is 10^-9^ per locus per cell division [12], and the copy number is not considered to carry substantial information regarding cell lineage membership in healthy tissue. Therefore, mutations are scarce when reconstructing lineage trees for healthy tissues, implying that more sophisticated models and computational methods are needed to capitalize fully on the existing mutations. This scarcity also highlights the need for single-cell DNA sequencing (scDNA-seq) data since it reveals more point mutations than any other current data type [13].

Regardless of its potential to reveal mutations, scDNA-seq data comes with its challenges [14–18]. Due to the small amount of genomic data available in a single cell, the genome needs to be amplified before sequencing [19]. Unfortunately, the whole-genome amplification methods, such as the multiple displacement amplification (MDA) method [20] and the multiple annealing and looping-based amplification cycles (MALBAC) method [21], introduce technical artifacts known as amplification errors (AEs) that are hard to distinguish from mutations. Moreover, so-called allelic dropout (ADO) events remain even after the amplification. In addition, the subsequent sequencing of the amplified materials introduces sequencing errors [22–25].

There have been several methods explicitly made for scDNA-seq data, both for identifying mutations (single nucleotide variant (SNV) callers) and for reconstructing cell lineage trees, although several of them are targeting cancer data. Monovar [26] is an SNV caller designed specifically for scDNA-seq data; for each position, it models the ADO with a Bernoulli distribution, the AEs with independent and identically distributed (i.i.d.) Bernoulli random variables and base-calling error probabilities depend on Phred quality scores, [27, 28], while utilizing dynamic programming. LiRA [29] and Conbase [30] are scDNA-seq SNV callers that leverage read-phasing, while the latter does variant calling based on the population of single cells.

There has been a sequence of single-cell tree reconstruction methods targeting cancer data, [31–36], leading up to the SCIΦ method [37]. Interestingly, for the cancer case, the infinite sites assumption (ISA, [38–40]) may be violated due to segmental deletions. However, for healthy tissue, the ISA is an appropriate assumption. So, since SCIΦ is based on ISA, it is also relevant to the analysis of healthy tissue. SCIΦ has a probabilistic model that allows joint SNV calling and tree reconstruction using the Markov chain Monte Carlo (MCMC) method. More recently, the Phylovar [41] method was shown to handle millions of loci and be faster than SCIΦ while having similar accuracy.

## Method overview

Scuphr is a distance-based phylogenetic inference method that reconstructs cell lineage trees from scDNA-seq data, produced by experimental procedures that amplify the cells’ genomes, using amplification methods such as the MDA method [20] and the MALBAC method [21]. Analyses of this data type need to distinguish somatic mutations from sequencing errors and nucleotide substitutions caused by amplification. Therefore, Scuphr relies on a probabilistic model of read-phasing and these two error sources. Read-phasing is a technique applied to identify which allele the read comes from, which is used to distinguish if the read could be from a mutated or a non-mutated segment. We first describe our model without read-phasing and then introduce the details of the read-phasing.

The amplification process is modeled as a generalized Pólya urn process, in which a drawn ball with an error probability is replaced by one of the same color and one of another, in contrast to the ubiquitous replacement by two of the same color in the Pólya urn. The observed Phred scores define the base-calling error probabilities. The model also contains a probability for the ADO events. Another vital part of Scuphr is a dynamic programming-based inference algorithm that, based on the error model, computes the probability that two cells have different genotypes at any investigated potential mutation site.

Scuphr processes the scDNA-seq data using a *site selection* method that identifies the candidate sites that will be analyzed using the probabilistic model and contribute to the distance. The distance is obtained by combining the probabilities of different genotypes across the selected sites for each pair of cells. Finally, this distance is used as an input to a distance-based phylogenetic method, the neighbor-joining (NJ) algorithm [42].

The summary of the Scuphr workflow is shown in Fig 1. The input to Scuphr consists of bulk and single-cell DNA reads. First, candidate mutation sites are detected using the site selection method, Fig 1a.These candidate mutation sites can consist of a single base pair, like in most state-of-the-art methods, or two base pairs where the candidate mutation site is accompanied by a nearby germline SNV (gSNV). We call these site types *singleton sites* and *paired sites*, respectively. These site types will be referred to as *sites* throughout the paper. Second, site-associated distance matrices are calculated in parallel for each selected site, Fig 1b, Eq 3. Third, a single distance matrix is obtained by combining the site-associated distance matrices, Fig 1c, Eq 1 and 2. Finally, the cell lineage tree is reconstructed by applying the NJ algorithm to the final distance matrix, Fig 1d.In addition, the site-associated distance matrices can be sampled with replacement several times to obtain bootstrap lineage tree samples, which could be used to get consensus trees and edge supports.

**Fig 1.**
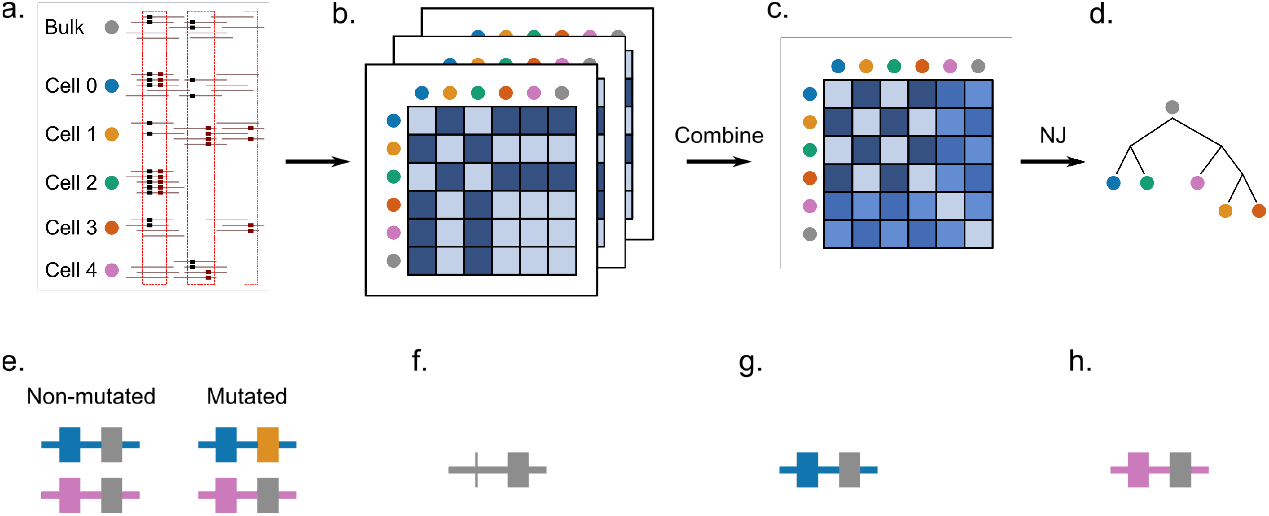
The workflow of Scuphr and the illustration of read-phasing. **a-d:** The workflow of Scuphr. **a:** The sites are selected for analysis from bulk and single-cell DNA sequencing data. **b:** The distance matrix of each chosen site is calculated independently. **c:** The distance matrices are combined. **d:** The cell lineage tree is constructed from the final distance matrix using the NJ algorithm. **e-h:** Illustration of read-phasing. The first base pair is the gSNV locus and the second base pair is the candidate site. The gSNV nucleotides are shown in blue and pink colors. The reference and the mutation nucleotides are shown in gray and yellow, respectively. **e:** The non-mutated and mutated genomes are displayed. The mutation is associated with the blue allele (blue gSNV nucleotide). **f:** An example read covering both sites, but the gSNV information is disregarded as in many methods of SNV calling and tree reconstruction. The reference nucleotide is observed at the candidate site. The read might belong to either the non-mutated or the mutated genome. **g:** An example read with available gSNV site information. We can *phase* the read (identify which allele it originates from). The read must come from the blue allele; in this case, the read comes from the non-mutated genome. **h:** An example read with available gSNV site information. The read must come from the pink allele; in this case, the read might belong to either the non-mutated or mutated genome.

The read-phasing assists in identifying missing data and separating somatic mutations and errors using patterns of co-occurrence of the nucleotides at the gSNV loci and the candidate sites. Fig 1e shows an example of a non-mutated and a mutated cell’s genome. Each cell has two alleles; one maternal and one paternal. The first locus is a gSNV, where the nucleotides at the first and second allele differ. The second locus is the candidate site, where the non-mutated cell has the reference nucleotide in both alleles, and the mutated cell has a reference and an alternate nucleotide. The non-mutated and mutated cells have the same second allele, and their difference is due to the mutation located at the first allele. The mutation is associated with the blue gSNV nucleotide. In Fig 1f, a read with the candidate site is shown. One cannot decide if this read comes from a non-mutated or a mutated genome since no information is available for the gSNV locus. Therefore, it cannot be attributed to any of the alleles. In Fig 1g, a read from the site pair is observed. Since both nucleotides are observed, and the blue gSNV nucleotide accompanies the candidate reference nucleotide, one can conclude that the read comes from the non-mutated genome. However, in Fig 1h, the reference nucleotide is accompanied by the pink gSNV nucleotide, which could come from either of the genomes.

## Results

The two state-of-the-art methods, SCIΦ [37] and Phylovar [41], can exploit amplified, diploid scDNA-seq data. According to the recently published paper, SCIΦ and Phylovar perform so similarly in terms of accuracy of SNV calling that it is hard to identify the one with the highest accuracy, while Phylovar takes less time to complete its analysis by taking advantage of the efficient vectorized computations [41]. We compared the performance of Scuphr with SCIΦ in terms of cell lineage tree reconstruction accuracy. Moreover, we also demonstrate the potential of Scuphr in terms of runtime; its runtime scales with the available number of cores, and we compare it with SCIΦ. This decision is motivated since, with a sufficient number of cores, Scuphr would be faster than Phylovar as well.

### Accuracy evaluation of synthetic datasets

To evaluate the performance of cell lineage tree reconstruction, we compared the topologies of the trees inferred by Scuphr and SCIΦ with the ground truth cell lineage tree. We use a similarity measure defined as one minus the normalized Robinson-Foulds (RF) distance.The similarity score is in [0, 1], where 1 means the tree topologies are the same. We investigated the accuracy across several combinations of AE rates, ADO rates, frequencies of loci with paired sites, and the number of cells. We investigated two AE rates: 10^-5^, see Fig 2 and 10^-3^, see Fig 3. First, for each choice of other parameters, the frequencies of loci with paired sites considered were 0.001, 0.01, 0.1, and 1. Second, for each choice of the other parameters, the ADO probabilities considered were 0, 0.1, and 0.2. Third, all parameter configurations were investigated for inputs having 10, 20, and 50 cells. Finally, Phred scores and read errors were generated. The details are presented in Methods.

**Fig 2.**
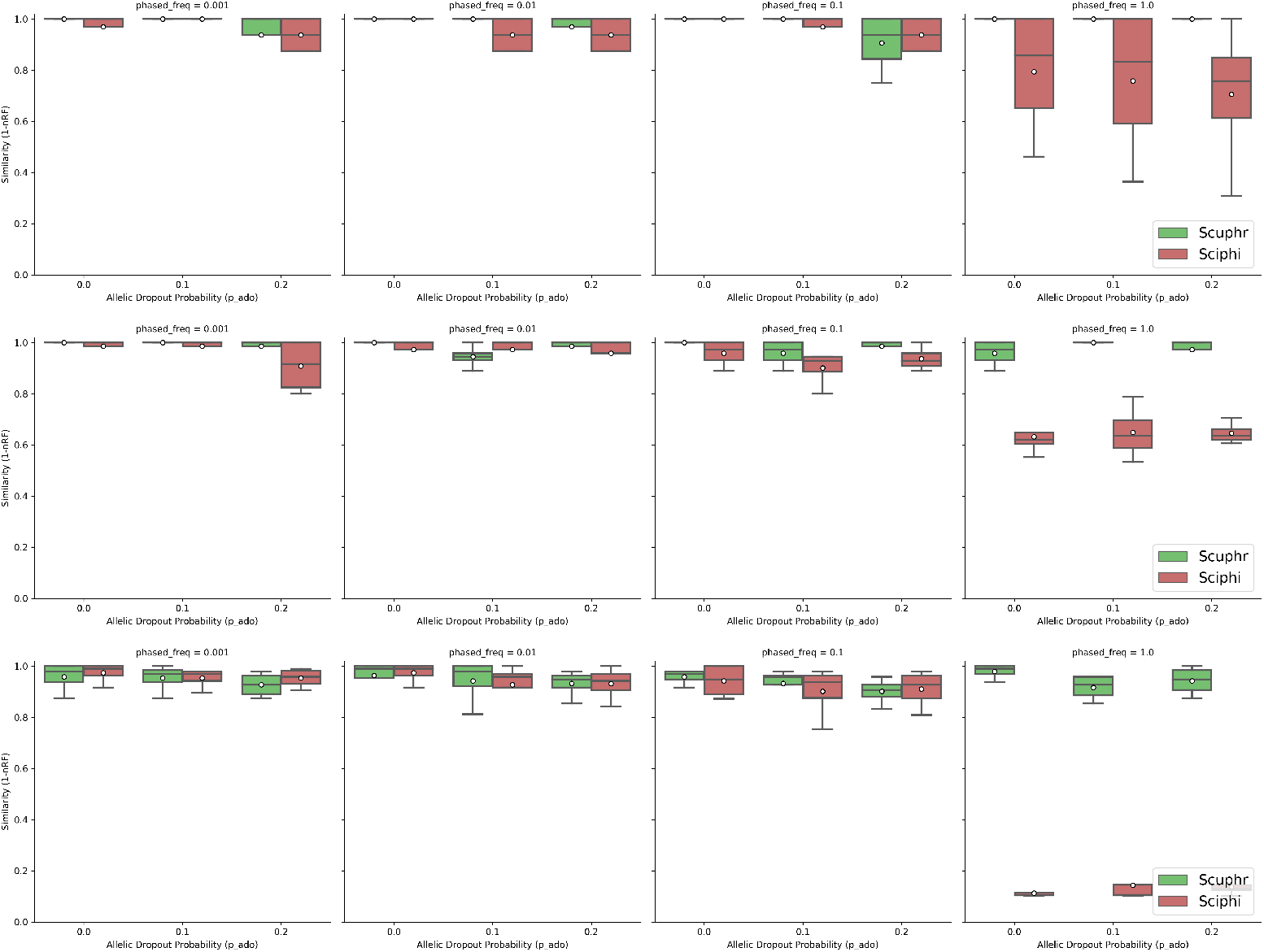
Similarities between the true and inferred trees for low amplification errors. **Top row:** Results for 10 cells. **Center row:** Results for 20 cells. **Bottom row:** Results for 50 cells.

**Fig 3.**
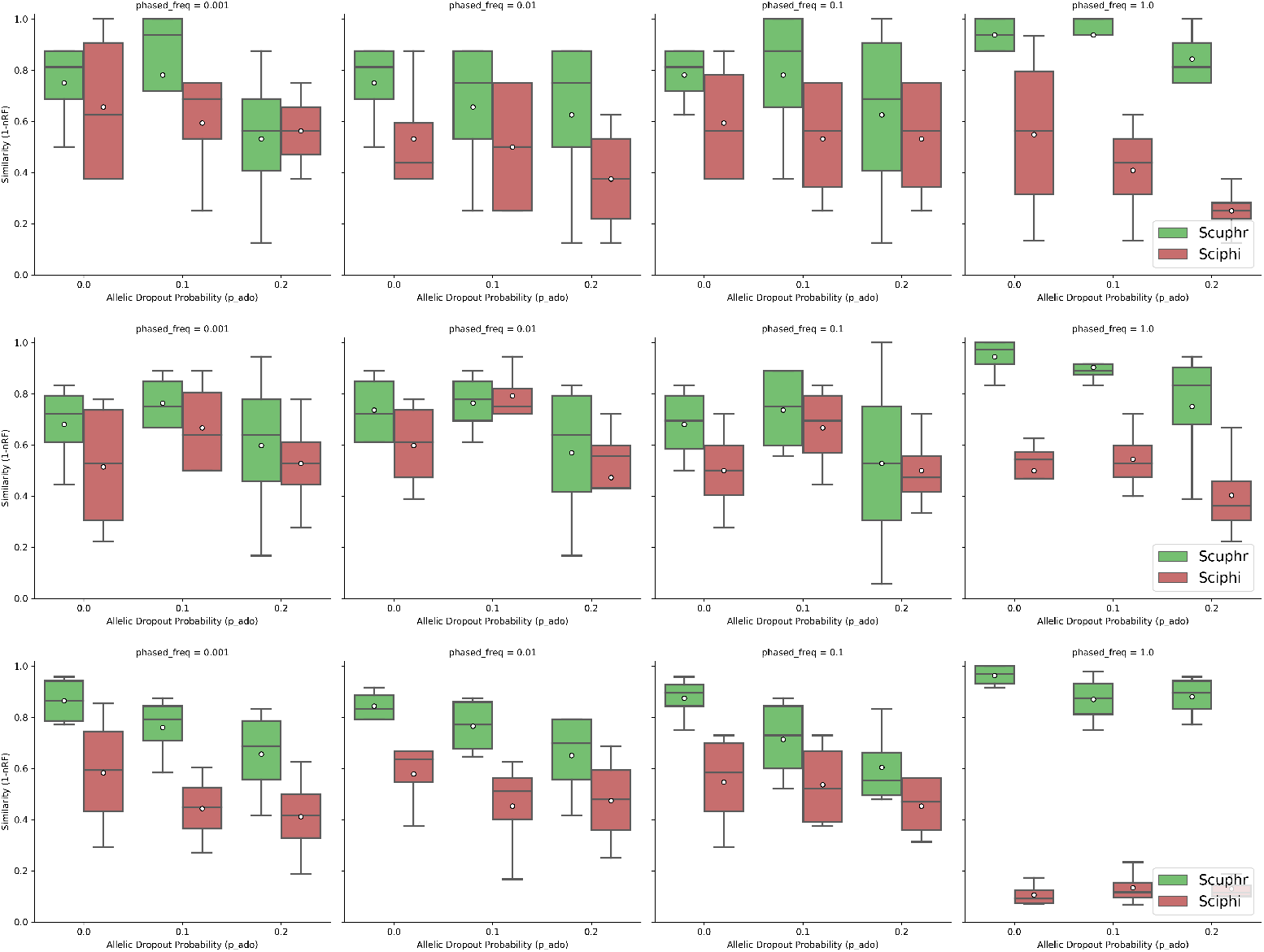
Similarities between the true and inferred trees for high amplification errors. **Top row:** Results for 10 cells. **Center row:** Results for 20 cells. **Bottom row:** Results for 50 cells.

Both methods perform well for the lower error rate 10^-5^, Fig 2. However, with a few exceptions, Scuphr has a higher mean accuracy, and other quantiles are higher for Scuphr than for SCIΦ. This trend is accentuated for the case where the frequency of loci with paired sites is 1.

For the higher error rate of 10^-3^, Fig 3 shows a readily noticeable difference between the two methods. The average accuracy for Scuphr is always better than that of SCIΦ, and other quantiles are, in almost all cases, higher for Scuphr than for SCIΦ. In particular, when all of the loci are paired, Scuphr exploits the paired sites for read-phasing, but SCIΦ has more or less the same average accuracy for lower frequencies of paired sites. In this case, the average accuracy of Scuphr is almost twice as high as that of SCIΦ.

Interestingly, when SCIΦ is provided with the sites selected by our candidate site selection method, its accuracy is improved in several cases. However, its accuracy is also, in many instances, decreased. Moreover, for the biological data, it is, due to the number of sites, that it takes an even longer time to run SCIΦ with the sites selected by our site selection method. Therefore, in Fig 2 and 3, we presented the results of the entire SCIΦ method, as described in [37]. The accuracy obtained when SCIΦ is provided with the site selected by our candidate site selection method is described in the S1 Appendix.

#### Runtime analysis on synthetic datasets

In addition to the lineage tree reconstruction accuracy, we also compared the wall-clock runtime of Scuphr to that of SCIΦ. All runtime experiments were performed on a single cluster node with 32 CPU cores, and each configuration was repeated ten times. As Scuphr can be used with default and estimated parameters, the runtime analysis for the parameter estimation step was performed separately, and the results are presented in the S1 Appendix. We ran both methods with the same sites to compare the runtimes. The time used to obtain these sites is excluded from the runtimes reported here. We run SCIΦ with both single and multiple cores. The runtimes were very similar across the number of cores, and in this section, the single-core runtimes are presented for SCIΦ. For the multiple core runs, we direct the reader to the S1 Appendix.

Fig 4 shows how the runtime changes across fractions of singleton sites and the number of cores. The left, center, and right subfigures display the runtime analysis for 10, 20, and 50 cells, respectively. Since each distance matrix is calculated independently, the main part of our proposed algorithm is embarrassingly parallel, i.e., the wall-clock runtime scales linearly with the number of available cores, as seen in the figure. Additionally, the algorithm’s runtime is linear in the number of sites. For 10 and 20 cells, our software infers the lineage tree faster than SCIΦ. Our software is faster for singleton sites and 50 cells when at least two cores are used for computation.

**Fig 4.**
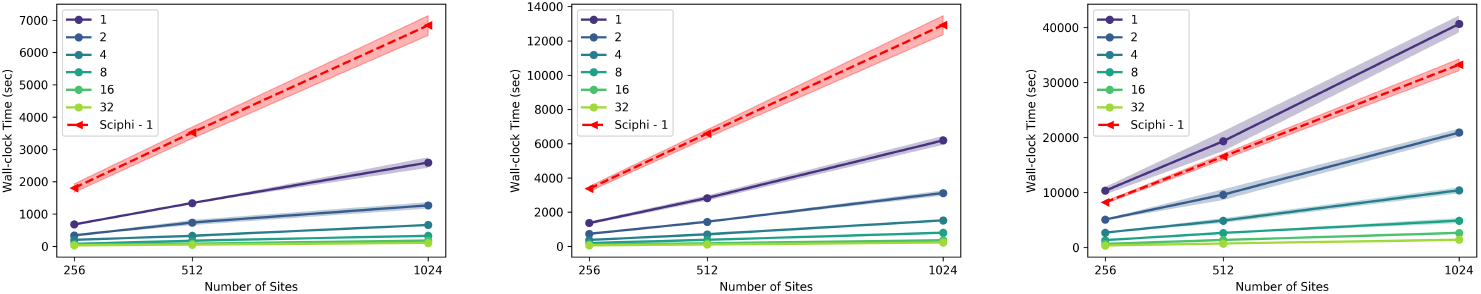
Runtime comparison for singleton sites. The x-axis is the number of sites, and the y-axis is the wall-clock time in seconds. The red dashed line is the runtime of SCIΦ using a single core. The remaining lines are the runtimes of the proposed method for varying numbers of cores. The left, center, and right subplots are the results for 10, 20, and 50 cells datasets, respectively.

The wall-clock runtime comparisons for paired sites are shown in Fig 5. Also, in this case, the left, center, and right subfigures correspond to 10, 20, and 50 cells, respectively. Our paired site analysis is slower than the singleton site analysis due to the number of fragment types considered. Nonetheless, the method has the same scalability trend as in the singleton site experiments. Our method is faster than SCIΦ for the 10 cells case when at least eight cores are used and for the 20 and 50 cells datasets when at least 16 cores are used.

**Fig 5.**
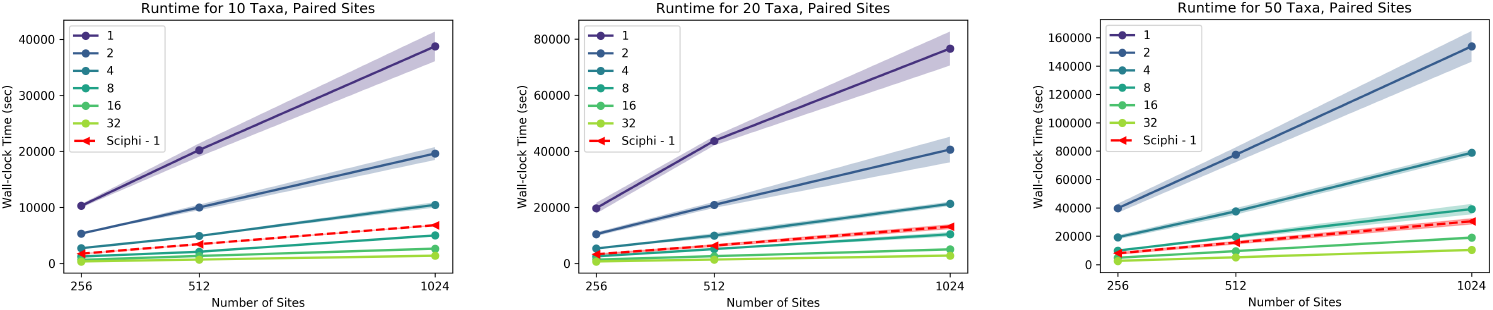
Runtime comparison for paired sites. The x-axis is the number of sites, and the y-axis is the wall-clock time in seconds. The red dashed line is the runtime of SCIΦ using a single core. The remaining lines are the runtimes of the proposed method for varying numbers of cores. The left, center, and right subplots are the results for 10, 20, and 50 cells datasets, respectively.

These runtime analyses were performed on a single cluster node. Nevertheless, one can use multiple nodes to compute the distance matrices without communication overhead. Consequently, Scuphr achieves a linear speedup in the total number of available cores on a cluster. The final step of Scuphr, the lineage tree reconstruction, runs on a single core; however, this step is very efficient and does not change the overall asymptotic runtime.

#### Biological data analysis

In this section, the accuracy of Scuphr and SCIΦ are compared using a fibroblast dataset previously used in [30].^1^ The dataset consists of scDNA-seq data for 18 cells with a recent common origin and a known lineage tree topology. This single-cell DNA data has been obtained by amplifying the DNA using MALBAC before the sequencing. These cells are so closely related that very few mutations distinguish them; hence, it is very hard to reconstruct the true lineage tree topology. The cells belong to two main monophyletic groups, one containing cells 0-11 and one containing cells 12-17. The dataset also includes a bulk DNA sample from the donor, which can be used as an outgroup.

The data were preprocessed using the pipeline described in S1 Appendix, and the sites of interest were identified as described in Methods. More than 3 million sites were selected for analysis; the details are presented in S1 Appendix.

Since the fibroblast dataset is so hard that a reconstruction method would at most identify the two main monophyletic groups correctly, we devised a test based on bootstrapping, using the transfer bootstrap expectation (TBE) [43] edge supports. 100 bootstrap lineage trees were constructed by bootstrapping sites (hence, bootstrapping the distance matrices). The TBE supports of the bootstrapped trees on the true lineage tree topology were computed with the Booster software [43]. The TBE support of a branch *b* is in the [0, 1] range where “0 means that the bootstrap trees contain the edge *b* in a random fashion and 1 means that *b* appears in all bootstrap trees” [43]. Scuphr separates the two main monophyletic groups with very high TBE support, 0.8, Fig 6a. Also, all branches within two smaller monophyletic groups, cells 4-5 and cells 10-11, respectively, are correctly inferred in all bootstrap rounds.

**Fig 6.**
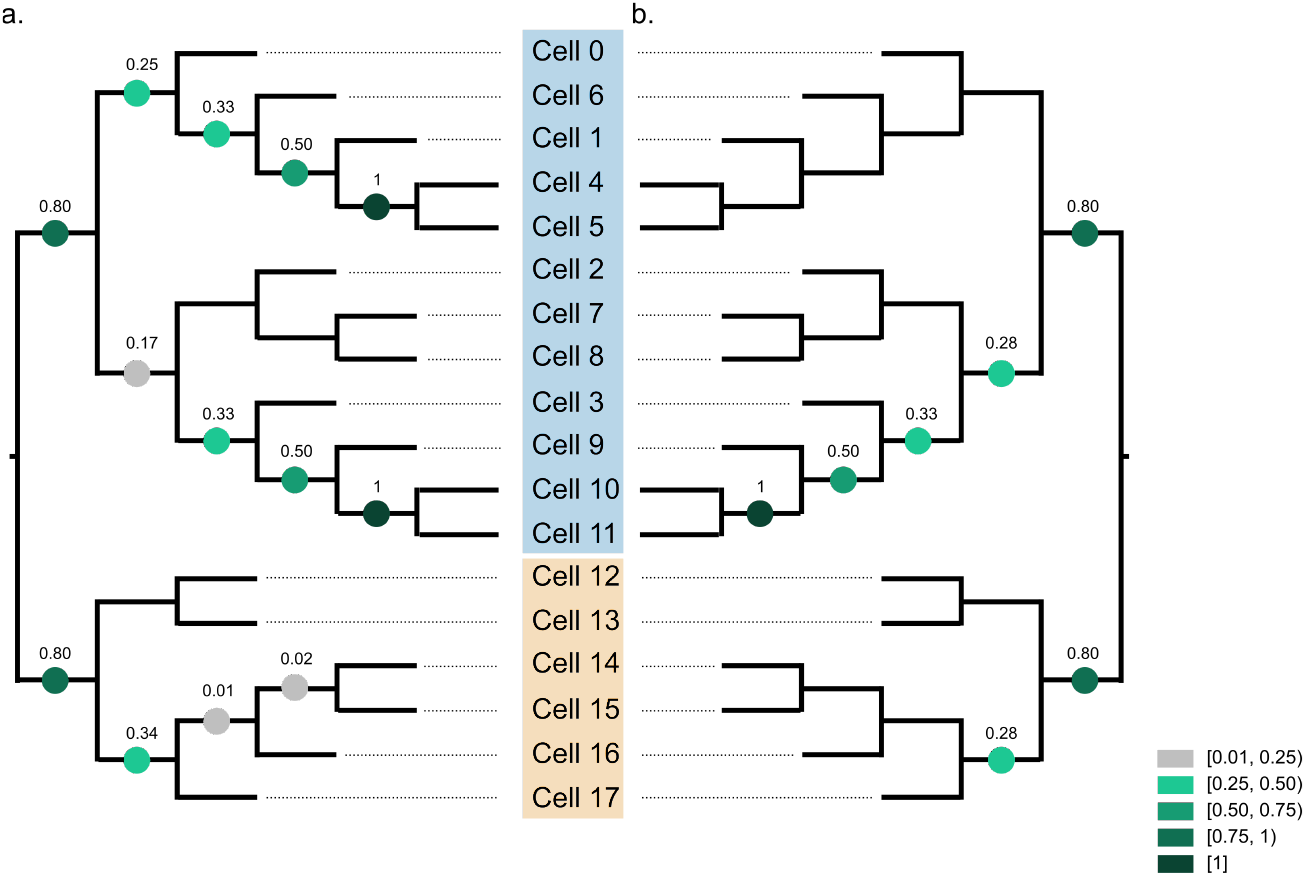
The TBE supports of bootstrap trees are projected onto true lineage tree topology. The clones are marked with blue and beige colors. **a:** TBE supports of Scuphr. **b:** TBE supports of SCIΦ.

To compare accuracy, we also applied SCIΦ to the fibroblast data. Due to the input format requirement (bulk and single-cell sequencing data in Mpileup format), we could not run SCIΦ on the whole genome in a single run. Instead, we applied SCIΦ independently to each chromosome, see S1 Appendix for details. We sampled 100 bootstrap trees from the lineage trees reported by SCIΦ for the chromosomes; the trees were weighted by the number of identified mutations per chromosome. The TBE supports are evaluated on the true lineage tree topology (Fig 6b). SCIΦ got the same TBE supports for a subset of branches (separation of two clones and the subclone consisting of cells 3, 9, 10, and 11). SCIΦ reported higher support for a single branch, 0.28, than Scuphr, 0.17. For the rest of the branches, Scuphr reported higher support or equal support to that reported by SCIΦ. In the monophyletic group consisting of cells 0, 1, 4, 5, and 6, in contrast to SCIΦ, Scuphr inferred substantial subclonal structure.

## Materials and methods

In this section, we describe the proposed model step-by-step. First, we present the probabilistic graphical model of Scuphr in detail and outline important components and formulas. Second, we describe how to reconstruct the cell lineage tree from the outcome of the first part. Third, we describe the candidate loci selection criteria in detail. Fourth, we show how the model parameters are estimated. Fifth, the simulated data generation procedure is presented. Finally, two accuracy metrics, the similarity score, and the TBE support, used in the study are described.

### The probabilistic model

First, we introduce some important concepts. Recall that, as most state-of-the-art methods do, a set of candidate mutation loci is used for analysis. We call a candidate locus that covers a single base pair a *singleton site*. Moreover, Scuphr can facilitate gSNVs near candidate loci and do read-phasing. A candidate mutation locus paired with a gSNV site is called a *paired site*. We will refer to all site types as *sites* throughout the Methods section for brevity. Unless otherwise stated, the model descriptions hold for all site types. Let Π be the set of sites selected for analysis and *C* be the number of single cells.

#### The probabilistic graphical model

Fig 7 shows the probabilistic graphical model of Scuphr at *π* ∈ Π. *a, b*, and *α* are the model hyperparameters. *p_ado_, p_ae_*, and *p_m_* are the model’s parameters and correspond to the ADO, AE, and mutation probabilities. A set of reads, their corresponding base-calling error probabilities, and the coverage of each cell *c* are observed and represented by **R***_c_*, **Q***_c_*, and *L_c_*. Moreover, the bulk genotype, *B*, is observed. The single-cell mutation status is represented with *G_c_; G_c_, B*, and the common mutation type random variable, *Z*, define the single-cell genotype, *X_c_*. The aforementioned scDNA-seq specific challenges are modeled with 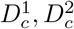, and *A_c_* random variables; 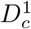 and 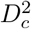 model the ADO events of each allele, and *A_c_* represents the number of errors that have happened during amplification. The fragment types generated at the end of the amplification process and their counts are expressed by *F_c_* and *N_c_*, respectively.

**Fig 7.**
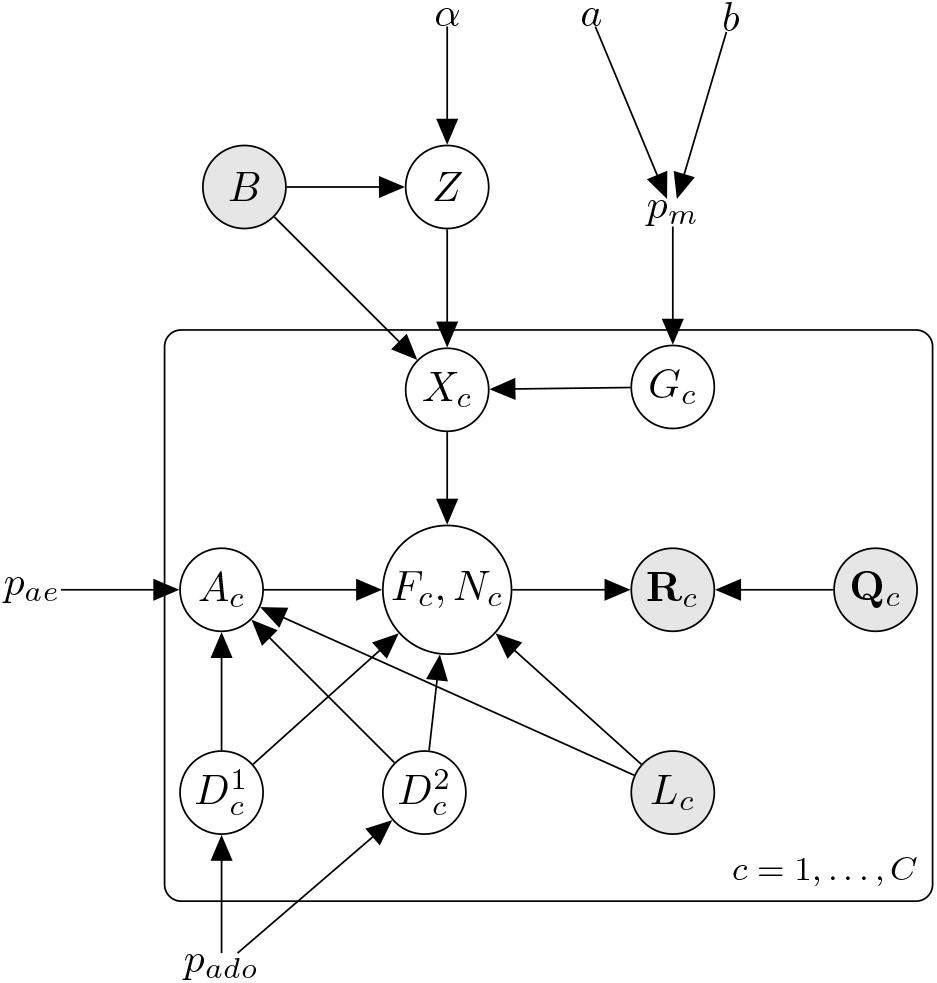
The graphical model of Scuphr for a site *π*. The shaded nodes are the observed random variables.

The graphical model consists of cell lineage, DNA amplification, and read sequencing.

- **Cell lineage:** At *π*, all the non-mutated cells have the bulk genotype, and all the mutated cells must share the same mutation type under the ISA. This shared mutation type is modeled with a Dirichlet-Categorical distribution with the hyperparameter *α*. The cells’ mutation statuses are modeled with i.i.d. Bernoulli random variables with a mutation probability, *p_m_*. The mutation probability has a Beta prior distribution with hyperparameters *a* and *b*. The mutation status random variable, bulk, and mutation type variables define the genotype of the cell, *X_c_*.
- **DNA amplification:** Here, we model the ADO events of each allele with Bernoulli random variables, with the same ADO probability *p_ado_*. The number of AEs that have happened during the amplification is modeled with a Binomial random variable with probability *p_ae_*, and its number of trials depends on the ADO random variables and the observed read coverage. These ADO and AE random variables, the observed read coverage, and the single-cell genotype form the amplified fragments, *F_c_* and their corresponding counts, *N_c_*.
- **Read sequencing:** Finally, the amplified fragments are sequenced and create observed reads. Since the read sequencing is an erroneous process, the observed Phred scores are used to obtain the base-calling error probabilities, *Q_c_*, and the uncertainty of read sequencing is modeled.

We briefly discussed the graphical model. More details of its components are presented in the following parts of this section.

#### Distance matrix

For each selected site, *π* ∈ Π, we construct a symmetric nonnegative (*C* + 1) × (*C* + 1) distance matrix *M^π^*. *M_c,c′_* is the distance between single-cells *c* and *c′*, computed by

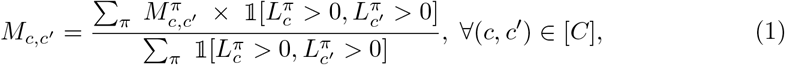

where 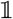 is the indicator function, and 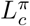 is the coverage of *c* at *π*. Only the sites where both cells have coverage are considered during distance computation. The matrix’s last row and column are the distances between the cell and the non-mutated bulk, *b*, computed by

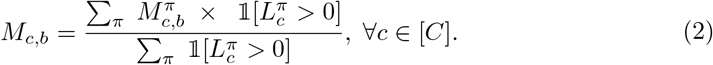

#### Distance between two single-cells at *π*

The binary random variable 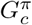 represents the mutation status of single-cell *c* at *π*: 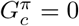 if the cell is not mutated and 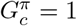 if the cell is mutated. The distance between two single-cells at *π* is

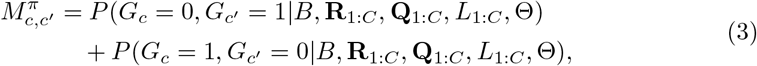

where *B* is the bulk genotype, Θ = {*α, a, b, p_ado_, p_ae_*} is the set of model parameters and hyperparameters, and **R**_1:*C*_, **Q**_1:*C*_, and *L*_1:*C*_ are the observed reads, base-calling error probabilities, and read coverages of single-cells.^2^ On the random variables of the right-hand side of the above equation, we omit the *π* superscript and use the same convention in the following parts.

The mutation status probability of two cells at a site satisfies

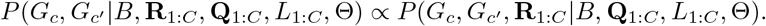

#### Lineage model

The ISA implies that (i) a site can be mutated at most once, and (ii) all the mutated cells at the loci share the same mutation. We marginalize the mutation types

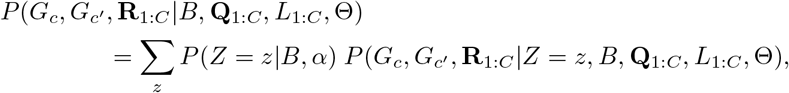

where *Z* is the mutation type random variable and follows the Dirichlet-Categorical distribution as described previously;

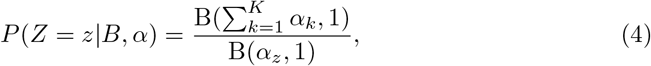

where *K* is the number of possible mutation genotypes, α is the concentration parameter, and *B* is the Beta function. *Z* differs from the non-mutated bulk genotype by a single nucleotide, e.g., we may have *B* = (*A, A*) and *Z* = (*A, G*) for a singleton site or *B* = (*AA, AT*) and *Z* = (*AA, GT*) for a paired site. See S1 Appendix for details.

#### Marginalization of other single-cells

Using the notation *G*_1:*C*\{*c,c′*}_ for the mutation status random variables of all cells except *c* and *c′*, the joint distribution of the reads and the mutation statuses of the single-cells *c* and *c′* can be expressed as

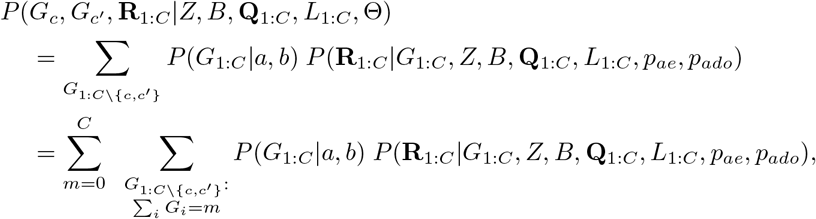

where 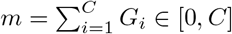 is the number of mutated cells at the site *π*. The summation over mutation counts and mutation statuses in the above equation can be computed efficiently using dynamic programming.

As mentioned earlier, we assign Beta prior distribution on the mutation probability, *p_m_*, with the hyperparameters *a* and *b*. Given *p_m_*, the mutation statuses of all single-cells are conditionally independent and are i.i.d. Bernoulli random variables. The joint distribution of single-cell mutation statuses is

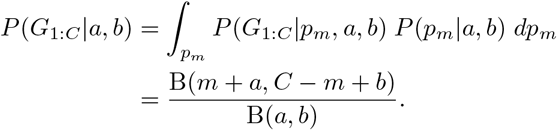

The derivation is presented in S1 Appendix.

#### Genotypes of single-cells

We define the auxiliary random variables, *X*_1:*C*_, to denote single-cell genotypes. The genotype of a single-cell *c* is

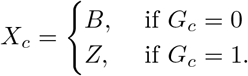

The values of *X*_1:*C*_ are deterministic functions of *G*_1:*C*_, *Z*, and *B*. From now on, we will use *X*_1:*C*_ notation instead of {*G*_1:*C*_, *Z, B*}, e.g.,

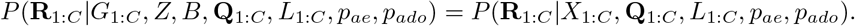

#### Conditional independence of single-cells

Given the genotypes of single cells, the amplification, and the allelic dropout probabilities, the likelihoods of reads are conditionally independent. The read likelihood is factorized by

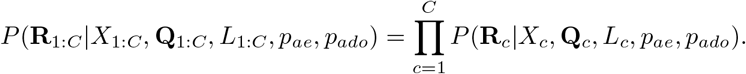

#### Introduction of fragments

We introduce the *fragments* created during the DNA amplification. The fragment types, *F_c_*, and their counts, *N_c_*, are marginalized as follows;

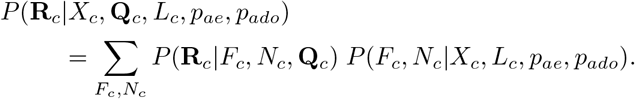

#### Amplification model

The DNA amplification is modeled with a generalized Pólya urn model. Two ADO events determine the initial state of the urn. These ADO events are modeled by two Bernoulli random variables with the same ADO probability, *p_ado_*;

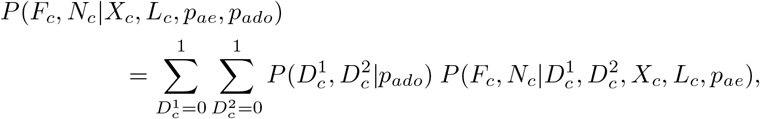

and

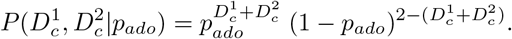

In the case of no ADO events, the process starts with one copy of each allele. The process starts with the other allele if there is one ADO event.

One can describe the urn process as follows; the urn is initialized with one or two colored balls. At each step, a ball is drawn from the urn, a copy of the ball is made, and both the original and the copy is put back into the urn. The outcome of this process can be represented by one or two lineage trees where the roots of the trees are the original copies of the alleles. We will refer to these trees as *amplification trees*. Given the initial state and the final number of balls in the urn, which is observed as the read coverages, the total number of edges in amplification trees is^3^

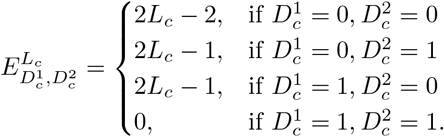

DNA amplification sometimes replaces a nucleotide; therefore, occasionally, the copy of a ball has a different color than the original. Let *A_c_* be the random variable describing the number of AEs has happened during the DNA amplification,

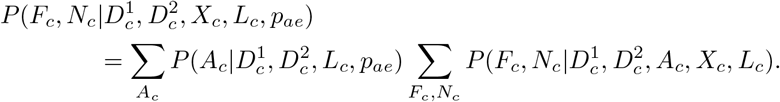

The probability of the number of AEs is a Binomial distribution over the edges of the amplification trees

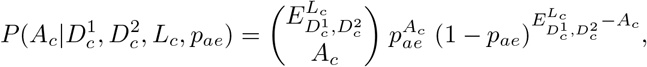

where *p_ae_* is the probability of an AE happening on an edge. In practice, *p_ae_* is very small (e.g., [10^-6^, 3 × 10^-4^] [19, 30]); hence we neglect the cases where *A_c_* > 1.

#### Marginalizing amplification trees

Let the fragment types and counts be 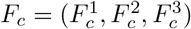 and 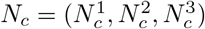, respectively. Let *d*(*F_i_*║*F_j_*) be the function that computes the Hamming distance between two fragment types, *F_i_* and *F_j_*.

The first two elements of *F_c_* are the cell genotype, 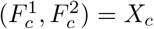, and the third element is the fragment type caused by an AE. In the case of no AE, 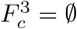. In the case of one AE, 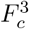 must differ a single nucleotide from its originating fragment, either 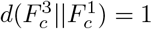 or 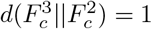.

Similar to the *F_c_* tuple, 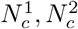, and 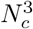 are the numbers of fragments of the first allele, second allele, and fragments carrying a nucleotide introduced by the AE. The total number of fragments is 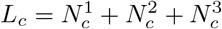. In case of an AE event, 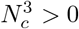; otherwise, 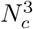 is zero. Finally, the first two elements of fragment counts must satisfy the ADO events, i.e., 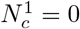 if 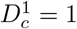 and 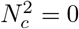 if 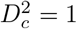.

With the above conditions satisfied, given the cell genotype, read coverage, ADO, and AE events, the probability of a fragment type and count pair is a product that contains up to three important factors. The first factor regards dividing *L_c_* fragments into one or two amplification trees. There is a single way to partition if there is an ADO event, e.g., (0, *L_c_*) if the first allele is dropped out or (*L_c_*, 0) if the second allele is dropped. Otherwise, due to the Polya urn, the number of reads follows a Beta-Binomial distribution, and each partition, {(1, *L_c_* – 1), (2, *L_c_* – 2),…, (*L_c_* – 1, 1)} has a 1/(*L_c_* – 1) probability. See S1 Appendix for details. The second factor regards the AE event; if an AE has happened, how many ways are there to get 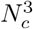 erroneous fragments? Here we should note that, even though we know how *L_c_* is partitioned into amplification trees, we do not know the internal structure of the trees. We need to consider all possible ways of forming the amplification trees (i.e., marginalization of the amplification tree topologies). We model an amplification tree as described in S1 Appendix. Assuming a count configuration of 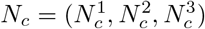 where 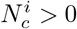 for all *i* ∈ {1, 2, 3} and that the AE is happening strictly on the first amplification tree, there are 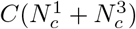 possible tree topologies of the first amplification tree, each with the probability of 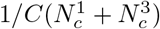. Moreover, 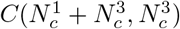 out of 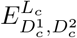 edges satisfy the specified count configuration in this marginalized amplification tree space. When all these are combined, the second component becomes 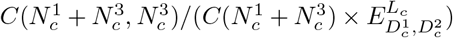. We direct the reader to Tables 1 and 2 in S1 Appendix for all the count and fragment type configurations. The final component is the probability of 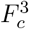 given AE; in the case of no AE, the probability is

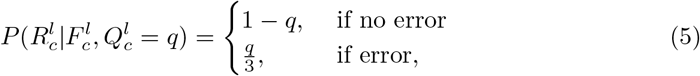

and in the case of AE, the probability is 1/3 for the singleton sites (1 × 3 = 3 different possible genotypes differ by 1 nucleotide), and 1/6 for paired sites (there are 2 × 3 = 6 possibilities).

The product of the three components leads to the fragment type and counts probability; 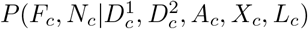, which is detailed in Tables 1 and 2 in S1 Appendix.

#### Read sequencing

Read sequencing is also erroneous and depends on sequencing technology [22–25]. We use the Phred quality scores (*ρ*) to compute the base-calling error probabilities, *Q*, [27, 28]; *Q* = 10^-0.1 × *ρ*^.

For a single read with a known originating fragment, the likelihood of the read at a singleton site is

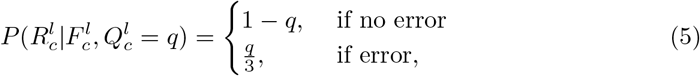

and the likelihood of the read at a paired site is

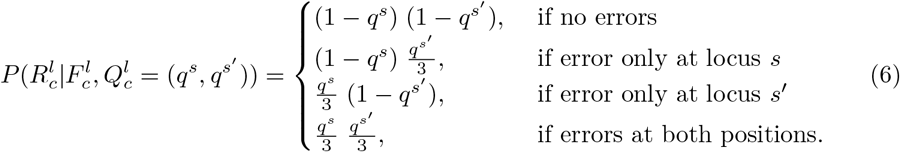

We use dynamic programming to efficiently compute the likelihood of multiple reads, 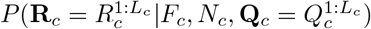, of cell *c*. The pseudocode is shown in S1 Appendix.

### Lineage tree reconstruction

The standard NJ algorithm [42] and its variants, such as FastNJ [44], are commonly used for distance-based methods. In the final step of Scuphr, the standard NJ algorithm is applied to the distance matrix to reconstruct the cell lineage tree using the implementation in the Dendropy library [45]. The tree is re-rooted, so the bulk node becomes the root and indicates the non-mutated state.

### Site selection

We use several heuristics to identify candidate loci for analysis. Even though Scuphr can run on the sites with no observed alternate nucleotides, these sites would not contribute information about the topology of the lineage tree and waste computational resources. Instead, we select a subset of the genome that could provide information regarding the topology.

#### Paired site selection

The main goal for the paired site selection is to find pairs of loci with a sufficient amount of alternative nucleotides *and* a heterozygous site nearby that can be used for read-phasing.

First, we identify the gSNVs using the unamplified bulk reads taken from another tissue. We run FreeBayes [46] software and set the read depth threshold to 10 and alternative nucleotide frequency to 0.2. The sites with heterozygous genotypes are considered the gSNV sites. Second, we check single-cell reads that cover the gSNV site and ensure at least two single-cells display the gSNV; that is, both nucleotides must be present in at least 20% of the reads from both cells. After this verification, we look for the candidate mutation sites around the gSNV. Both gSNV and the candidate sites must be covered with the same read to facilitate read-phasing.^4^ The reference nucleotide of the candidate site is determined from bulk reads; the site must have at least 10 reads in bulk data, and at least 80% of the reads are one nucleotide, which is referred to as the *reference*. For a site to be picked, at least 2 and at most *C* – 1 cells must agree on an alternative nucleotide (at least 20% of the reads should be different from the reference).^5^ If multiple gSNVs are near the candidate site, the closest gSNV is used to form the pair. Finally, one last gSNV check is done to ensure the single-cell reads (that cover both the candidate and gSNV sites) meet the gSNV requirement described above.^6^

#### Singleton site selection

During the singleton site selection, the gSNV heuristics are omitted. The candidate mutation site identification is performed using the same heuristics as the paired site selection.

#### Hybrid site selection

In the hybrid case, the algorithm works with both paired and singleton sites. A candidate mutation site is paired with a gSNV if the paired site criteria are met; otherwise, the site is picked as a singleton site.

### Parameter estimation and hyperparameter settings

We run the Metropolis-Hastings algorithm for 5, 000 iterations with three different initial values to infer the parameters *p_ado_* and *p_ae_*. We discard the first 20% samples as burn-in. The acceptance ratio of our Metropolis-Hastings algorithm is

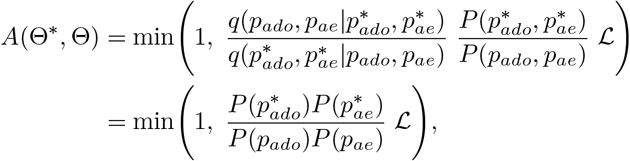

where

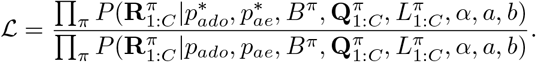

The likelihood is calculated similarly to the earlier derivations in Methods;

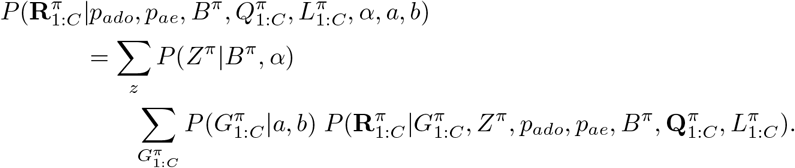

We use a Gaussian random walk proposal in which each parameter is treated independently, and samples are proposed using a Gaussian distribution with a 0.01 standard deviation. We set uniform prior probabilities for the parameters, calculate the likelihood of the observed reads 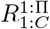 based on our model, and accept or reject the samples. The means of the samples are used as *p_ado_* and *p_ae_* parameters during the analysis.

Scuphr has three hyperparameters; *α, a*, and *b*. *α* is the concentration parameter of the Dirichlet-Categorical distribution used for the mutation type probabilities. We set *α* to all-ones vector. The *a* and *b* hyperparameters are for the Beta prior to mutation probability *p_m_*. We assigned uniform prior to the mutation probability by setting *a* =1 and *b* = 1. However, the user can set these parameters and, thereby, modify the algorithm’s tendency towards mutations.

For our experiments, we randomly picked 20 sites that are used to estimate the parameters. We sampled the initial value of *p_ado_* from *U*[0, 1] and *p_ae_* from *U*[0, 0.1]. We set the initial range of *p_ae_* to [0, 0.1] because the AE probabilities are reported to be small [19, 30].

### Simulation of synthetic datasets

We generated the synthetic dataset as follows. First, we generated random binary cell lineage trees with *C* leaves. We assigned 2 × (*C* – 1) × *μ* mutations, *μ* ∈ {10, 20}, to the tree’s edges and ensured each edge had at least one mutation. For each dataset, we generated a 1 million base pairs long diploid genome that was used for the bulk and the single cells. We randomly picked mutation loci from odd-numbered bases in the genome. For each of the phasing frequencies *ρ* ∈ {0.001, 0.01, 0.1, 1}, we picked *ρ* × 500, 000 base pairs as gSNV sites and placed them randomly in even indexed locations in the genome. So for *ρ* = 1, every second position in the genome is a gSNV site, and each read containing a mutation has an accompanying gSNV site. This construction is sufficient since the distance between sites does not affect the site selection or the subsequent analysis. The gSNV sites are shared by the single-cell genomes, and the mutation sites are shared according to their placement in the cell lineage tree. Further, we masked the single-cell genomes to account for cell-specific ADO events. For each pair of sites (which consists of consecutive positions, one even and one odd), we dropped the maternal and paternal alleles independently with *p_ado_* ∈ {0, 0.1,0.2}.

The fragments were generated using the Pólya urn process for each site. The masked single-cell genome determined the initial state of the urn. If both alleles are dropped out, no fragments were generated. In the case of a single ADO, all the fragments are generated from the non-dropped allele. If there is no dropout, the fragments are simulated from both alleles, i.e., the urn was initialized with two balls of different colors. The number of fragments per pair of sites was sampled from a Poisson distribution with rate parameter *λ* ∈ {10, 20}, i.e., an interval that contains the read depth found in our biological data. Whenever a fragment is copied, an AE occurs with *p_ae_* independently. So, even though we base our inference on assuming that there is at most one AE per site, we allowed multiple errors during the data simulation. We use *p_ae_* = 10^-5^ or *p_ae_* = 10^-3^ for all cells of a dataset; throughout the paper, we refer to these datasets as a dataset with *low* and *high* AEs, respectively.

We simulated how the fragments are sequenced so that reads are obtained, as follows. Phred quality scores were sampled from a discrete Uniform distribution in the range [30, 42]. A sequencing error was introduced based on the corresponding base-calling error probability.

We used a straightforward approach for the bulk reads and replicated the bulk genome 15 times. One can view this step as the bulk data is generated using 15 single-cell genomes without any mutations. This replication does not contain a DNA amplification step since the bulk data consists of reads sequenced from unamplified fragments.

### Accuracy metrics

This section describes the two accuracy metrics used for the analysis.

#### The similarity score

The Robinson-Foulds (RF) distance [47] is a symmetric difference metric commonly used for phylogenetic tree comparisons [36, 48]. The metric calculates the total number of bipartitions in either of the trees but not in both. We normalized the RF scores (nRF) by the total number of non-trivial bipartitions in both tree topologies,^7^ nRF = RF*/I_B_*, where *I_B_* is the total number of internal edges in the two tree topologies. Notice that the number of non-trivial edges of a tree depends on its topology, e.g., whether the tree is binary or not. We used a similarity score,

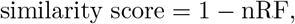

in [0, 1]. If the trees have the same topology, the similarity score is 1. If the trees do not have any common non-trivial bipartition, the similarity score is 0.

#### Transfer bootstrap expectation

The TBE is introduced as an alternative metric to compute the support of bootstrap trees on a reference tree topology [43]. Compared to Felsenstein’s bootstrap proportions [49], which checks how frequently an edge appears in the bootstrap trees, the TBE metric penalizes the slight topological differences less. TBE metric computes the number of operations (e.g., taxa removal) required to match an edge in the bootstrap tree to the reference tree.

TBE score of an edge is in [0, 1], where 1 means the edge exists in all bootstrap trees and 0 means the edge appears at random. The higher the TBE score of an edge, the better.

## Discussion

We evaluated the performance of Scuphr on a biological dataset and several simulated datasets and compared it with SCIΦ. Our investigation focused on the algorithm’s robustness for varying numbers of single-cells, read coverages, and technological artifacts such as AEs, ADOs, and sequencing errors. We observed that for the low amplification error datasets, Scuphr performs on par or better than SCIΦ in most cases.

For the high amplification error datasets, Scuphr consistently outperforms SCIΦ. When provided by the candidate mutation sites selected by our method, SCIΦ’s performance becomes similar to Scuphr in most low amplification error datasets; however, Scuphr continues to outperform SCIΦ in the high amplification error datasets.

In addition, we showed that the algorithm scales with the number of single cells and sites. Moreover, it scales inversely with the number of cores. For instance, using a single core, the main part of Scuphr takes approximately 1.6 hours for 20 cells and 1024 singleton sites. The time required for 100, 000 sites would be approximately 166.7 hours; however, with a modest infrastructure of five compute nodes with 32 CPU cores each, Scuphr’s runtime can be reduced to approximately 1 hour. This advantage makes it possible to analyze millions of sites in the genome, whereas most state-of-the-art methods can handle only a few thousand sites.

Finally, we evaluated the performance of Scuphr using a biological dataset of 18 single cells acquired from [30]. We selected approximately 3.4 million candidate sites for analysis and used bootstrapping to obtain edge supports on the reference tree topology. Although the biological dataset was challenging, Scuphr assigned high support values to the edges that separate the two main clones and some closely related cells. Additionally, Scuphr outperforms SCIΦ by obtaining higher edge supports for a subclone.

## Conclusion

Single-cell DNA sequencing technologies enable detailed analyses of development and cell differentiation [1–3]. We presented Scuphr, a probabilistic framework that reconstructs cell lineage trees from healthy, diploid single-cells using whole-genome amplified DNA sequencing data. Scuphr is designed with the challenges of the scDNA-seq data in mind, it fits well with the biological findings, and in particular, it obtains better accuracy with leveraging read-phasing.

In addition to the distance-based and MCMC-based methods, various variational inference based methods have recently been developed for tree reconstruction tasks [50–53]. These methods typically operate in the standard phylogeny setting and require a good set of initial trees for their analysis. In the potential future development of such methods where the domain moves toward the single-cell setting, Scuphr could provide a good set of bootstrap trees as input quickly.

Scuphr is designed for healthy, diploid scDNA-seq data. However, it can be enhanced to handle cancer data by incorporating copy number variations into its model. We will investigate how the extended model handles the challenges of single-cell tumor data and compare its performance with state-of-the-art methods in our future work.

## Supporting information

**S1 Appendix. Supplementary information.** The file includes additional formulations, biological dataset, and benchmarking details.

## Acknowledgments

Funding is provided by the Swedish Foundation for Strategic Research Grant BD15-0043. The computations and data handling were enabled by resources provided by the Swedish National Infrastructure for Computing (SNIC), partially funded by the Swedish Research Council through grant agreement no. 2018-05973.

## Footnotes

1 The dataset used in this study is a slightly modified version of the dataset in [30]. For details, see S1 Appendix.

2 The distance to non-mutated bulk is simply 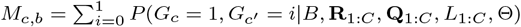

3 An additional incoming edge to the root is introduced to account for subsampling.

4 All nucleotides covered by a read come from the same allele.

5 The signal from a single-cell or all single-cells does not contribute to the information for lineage tree reconstruction.

6 The further the candidate site is from the gSNV site, the fewer reads cover both sites.

7 A leaf’s edge is considered trivial. If two tree topologies have the same leaf set, there will be edges that define the same bipartition. There will be *C* identical bipartitions, regardless of the internal structure of the trees.

