## Appendix for "Scuphr: A probabilistic framework for cell lineage tree reconstruction"

### Contents

**A Illustration of random variables** **2**

**B Dynamic programming algorithm to compute read probabilities** **3**

**C Common mutation type probability** **4**

**D Mutation status probability** **5**

**E Pólya urn model** **6**

**F Details on counting amplification trees and edges** **7**
F.1 $C(t)$ **derivation** . . . . . 8
F.2 $C(t, d)$ **derivation** . . . . . 8

**G Read likelihood** **9**

**H Differences between singleton and paired sites** **12**

**I Real data preprocessing** **13**

**J Fibroblast dataset information** **14**

**K Number of sites in biological data experiments** **15**

**L Similarity score comparison of all methods** **16**

**M Runtime analysis for parameter estimation** **18**

**N Runtime comparison with more cores for SCIΦ** **19**

**O SCIΦ details** **20**

#### A Illustration of random variables

Fig 1 shows the cell lineage, DNA amplification, and sequencing steps described in Methods. For simplicity, the site superscript $\pi$ is omitted.

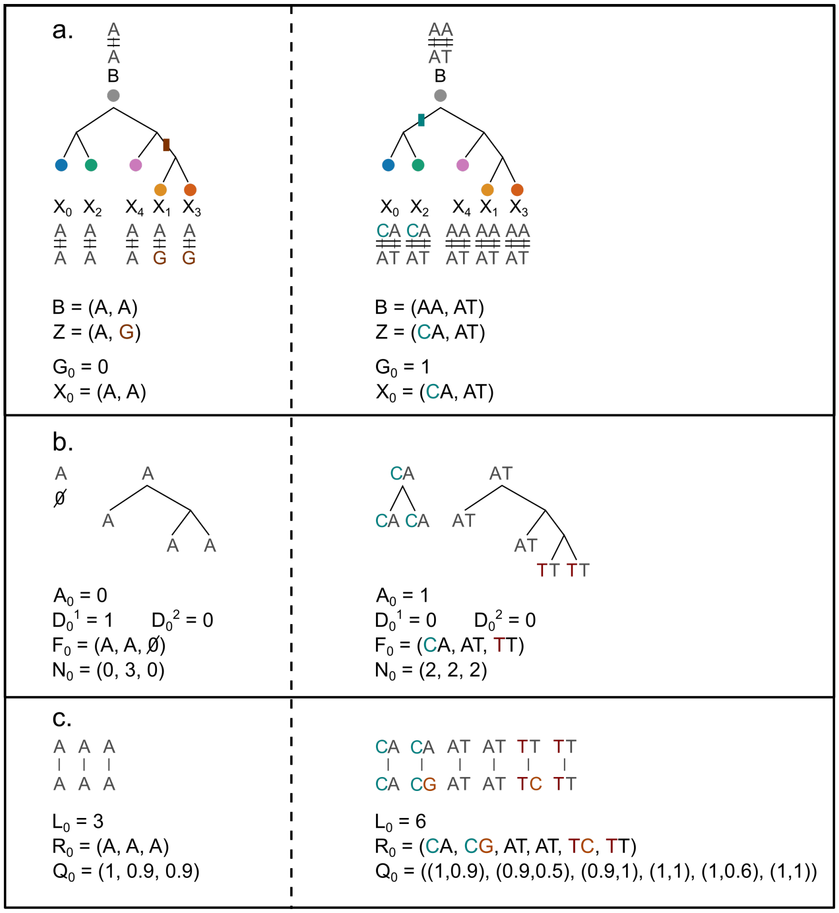


**Fig 1. Illustration of the random variables for a singleton site on the left column and a paired site on the right column. a:** The cell lineage tree. Bulk, common mutation type, and single-cell genotypes for the sites of interest are shown. Mutations and the branch they originate from are colored differently. **b:** The DNA amplification step for cell 0 is illustrated. AE and ADO event indicators are specified. The corresponding fragment types and the counts are written. **c:** The sequencing step is illustrated; the reads and the base-calling error probabilities are generated from the fragments. Sequencing errors are colored differently.

### B Dynamic programming algorithm to compute read probabilities

Algorithm 1 shows the precomputation of the read probabilities of a single cell given the corresponding base-calling error probabilities and fragment types. The probability of a single read, $P(R^l|Q^l, F^1)$, is computed as shown in Eq 5 and 6 in the main manuscript. The read probability given the fragments and the base-calling error probabilities, $P(\mathbf{R}_c|F_c, N_c, \mathbf{Q}_c)$, is obtained from the corresponding element in the $T$ table, $T[N_c^1, N_c^2, N_c^3]$.

---

**Algorithm 1** Dynamic programming algorithm to compute read probabilities

---

**Input:** Reads $\mathbf{R}_c = \{R_c^1, \dots, R_c^{L_c}\}$, error probabilities $\mathbf{Q}_c = \{Q_c^1, \dots, Q_c^{L_c}\}$ and fragment types $F_c = (F_c^1, F_c^2, F_c^3)$.
**Output:** $P(\mathbf{R}_c|F_c, N_c, \mathbf{Q}_c)$

**function** $retrieve(T, i, j, k)$:
  **if** $i < 0$ **or** $j < 0$ **or** $k < 0$ **then**
    **return** 0
  **else if** $i = j = k = 0$ **then**
    **return** 1
  **else**
    **return** $T[i, j, k]$
  **end if**

**function** $precompute(\mathbf{R}^{1,\dots,L}, \mathbf{Q}^{1,\dots,L}, F)$:
  allocate $T$
  **for** $i = 0, \dots, L$ **do**
    **for** $j = 0, \dots, L - i$ **do**
      **for** $k = 1, \dots, L - i - j$ **do**
        $l = i + j + k$
        $p_1 = P(R^l|Q^l, F^1) \times retrieve(T, i - 1, j, k)$
        $p_2 = P(R^l|Q^l, F^2) \times retrieve(T, i, j - 1, k)$
        $p_3 = P(R^l|Q^l, F^3) \times retrieve(T, i, j, k - 1)$
        $T[i, j, k] = p_1 + p_2 + p_3$
      **end for**
    **end for**
  **end for**
  **return** $T$

---

### C Common mutation type probability

The probability mass function of a Dirichlet-Multinomial distribution is

$$P(\mathbf{x}|n, \alpha) = \frac{n\ \mathrm{B}(\sum_{k=1}^{K} \alpha_k, n)}{\prod_{k:x_k>0} x_k\ \mathrm{B}(\alpha_k, x_k)},$$

where $K$ is the number of categories, $n$ is the number of trials, the $\alpha$ vector is the Dirichlet concentration parameter, $x_{1:K}$ are the number of outcomes of each category satisfying $\sum_{k=1}^{K} x_k = n$, and B is the Beta function.

The common mutation type probability follows the Dirichlet-Categorical distribution, that is, a Dirichlet-Multinomial distribution with a single trial ($n = 1$). The probability of a common mutation type is

$$P(Z = z|B, \alpha) = \frac{\mathrm{B}(\sum_{k=1}^{K} \alpha_k, 1)}{\mathrm{B}(\alpha_z, 1)},$$

where $B$ is the bulk genotype, and B is the Beta function.

If $\alpha$ is a one-vector, the common mutation type probability becomes a discrete uniform probability over mutation types;

$$\begin{aligned} P(Z = z|B, \alpha) &= \frac{\mathrm{B}(\sum_{k=1}^{K} \alpha_k, 1)}{\mathrm{B}(\alpha_z, 1)} \\ &= \frac{\mathrm{B}(K, 1)}{\mathrm{B}(1, 1)} \\ &= \frac{(K-1)!\ 0!}{K!}\ \frac{1!}{0!\ 0!} \\ &= \frac{1}{K}. \end{aligned}$$

### D Mutation status probability

The single-cell mutation probability, $p_m$, follows a Beta distribution with hyperparameters $a$ and $b$. The cells' mutation status configuration probability follows $C$ Bernoulli trials with $m$ successes with probability $p_m$. The mutation status probability with $m$ mutations is

$$\begin{aligned}
P(G_{1:C}|a,b) &= \int_{p_m} P(G_{1:C}|p_m,a,b)\ P(p_m|a,b)\ dp_m \\
&= \int_{p_m} p_m^m (1-p_m)^{C-m}\ \frac{p_m^{a-1}(1-p_m)^{b-1}}{\mathrm{B}(a,b)}\ dp_m \\
&= \frac{1}{\mathrm{B}(a,b)} \int_{p_m} p_m^{m+a-1}(1-p_m)^{C-m+b-1}\ dp_m \\
&= \frac{1}{\mathrm{B}(a,b)}\ \frac{\Gamma(m+a)\Gamma(C-m+b)}{\Gamma(m+a+C-m+b)} \\
&= \frac{\mathrm{B}(m+a, C-m+b)}{\mathrm{B}(a,b)},
\end{aligned}$$

where B is the Beta function, $\Gamma$ is the Gamma function, and $m = \sum_{c=1}^{C} G_c$ is the number of mutated cells.

#### E Pólya urn model

We modeled the amplification as a Pólya urn model. For simplicity, we imagine maternal and paternal alleles as colored balls, red and blue. The ball and its copy are added to the urn whenever a ball is drawn. The goal is to find the distribution of colored balls when they reach a target size (in our case, the total number of reads, $L_c^\pi = r + b$).

$$
\begin{aligned}
P(R = r, B = b) &= \\
&= \binom{r-1+b-1}{r-1} \times \left(\frac{1}{2}\frac{2}{3}\cdots\frac{r-1}{r}\right) \times \left(\frac{1}{r+1}\frac{2}{r+2}\cdots\frac{b-1}{r+b-1}\right) \\
&= \binom{r+b-2}{r-1}\frac{(r-1)!(b-1)!}{(r+b-1)!} \\
&= \frac{(r+b-2)!}{\cancel{(r-1)!(b-1)!}}\frac{\cancel{(r-1)!(b-1)!}}{(r+b-1)!} \\
&= \frac{1}{r+b-1}.
\end{aligned} \tag{1}
$$

In the first line of the above derivation, the first term ($\binom{r+b-2}{r-1}$) represents all combinations of selected balls (i.e., BBRRB, RBRBB, RRBBB, ...). In order to reach $r$ red and $b$ blue balls at the end, one needs to select $r - 1$ red and $b - 1$ blue balls. The second term ($\frac{(r-1)!}{r!}$) represents the probability of selecting all first $r - 1$ balls red. The third term represents selecting all remaining balls with the color blue.

Equation 1 indicates that we cannot say anything specific about the distribution of $r$ and $b$ values; each setup has an equal probability, depending on the total number of colored balls $L_c^\pi$.

The described urn model is simply a Beta-Binomial distribution;

$$
\begin{aligned}
P(R = r, B = b) &= BB(n = r + b - 2, \alpha = 1, \beta = 1) \\
&= \binom{n}{r-\alpha}\frac{\mathrm{B}(r,b)}{\mathrm{B}(\alpha,\beta)} \\
&= \binom{n}{r-\alpha}\frac{(r-1)!(b-1)!}{(r+b-1)!}\frac{(\alpha+\beta-1)!}{(\alpha-1)!(\beta-1)!} \\
&= \binom{r+b-2}{r-1}\frac{(r-1)!(b-1)!}{(r+b-1)!} \\
&= \frac{1}{r+b-1}.
\end{aligned}
$$

In the case of an ADO event, the urn is initialized with a single ball (e.g., red) and $P(R = L_c^\pi) = 1$.

### F Details on counting amplification trees and edges

We model the amplification process of each allele as follows. Let the *t-tree* be a rooted binary tree with $t$ leaves in which the inner vertices are labeled from 1 to $t-1$, where the labels indicate the order of amplification. A *d-edge* is an edge that has $d$ leaves under it.

In order to form a $t$-tree, there is $\binom{1}{1}$ choice for the first amplification event (at the root), $\binom{2}{1}$ possibilities for the second event, and so on, which leads to the number of $t$-trees

$$C(t) = \begin{cases} (t-1)!, & \text{if } t \geq 1 \\ 0, & \text{otherwise.} \end{cases}$$

The number of $d$-edges in all $t$-trees is

$$C(t,d) = \begin{cases} \frac{2t!}{d(d+1)}, & \text{if } 0 < d < t \\ C(t), & \text{if } d = t \\ 0, & \text{otherwise.} \end{cases}$$

Our model considers the amplified fragments' subsampling during the read sequencing. For this purpose, we introduced an arbitrary incoming edge to the root node, which enables the $C(t,t)$ computation. Fig 2 shows all possible 4-trees and illustrates 3-edges in red and 4-edges in *dashed* format.

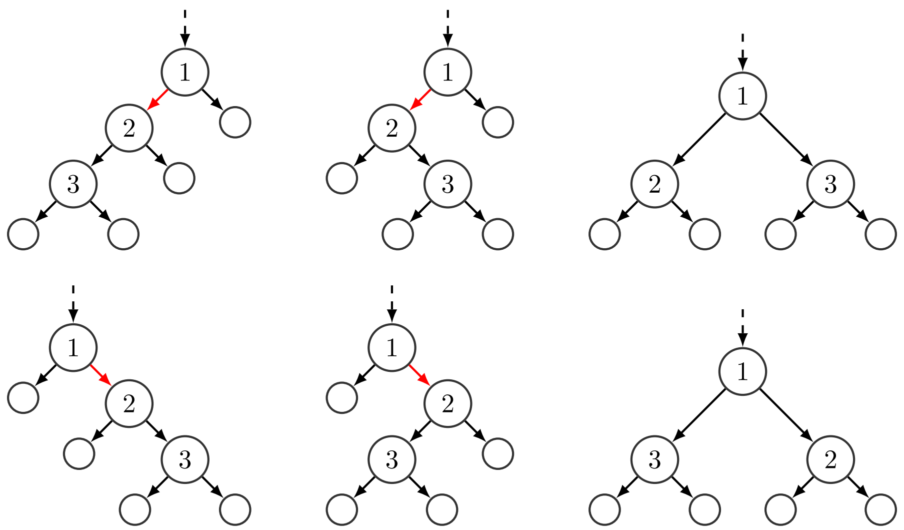

**Fig 2. Illustration of $C(4) = 6$ possible 4-trees.** The labeled nodes indicate the order of the amplification events. The dashed line represents an incoming edge to the root to account for subsampling during the sequencing of the fragments. The red edges are all possible 3-edges in 4-trees.

##### F.1 $C(t)$ derivation

The number of $t$-trees where $t \geq 1$ is computed as follows;

$$\begin{aligned}
C(t) &= \sum_{i=1}^{t-1} \binom{t-1-1}{i-1} C(i) \binom{t-i-1}{t-i-1} C(t-i) \\
&= \sum_{i=1}^{t-1} \binom{t-1-1}{i-1} C(i) C(t-i) \\
&= \sum_{i=1}^{t-1} \binom{t-2}{i-1} C(i) C(t-i) \\
&= \begin{cases} (t-1)!, & \text{if } t \geq 1 \\ 0, & \text{otherwise.} \end{cases}
\end{aligned}$$

##### F.2 $C(t,d)$ derivation

The number of $d$-edges in $t$-trees where $d < t$ is computed as follows;

$$\begin{aligned}
C(t,d) &= 2 \sum_{i=d}^{t-1} \binom{t-1-1}{i-1} C(i,d) C(t-i) \\
&= 2 \sum_{i=d}^{t-1} \frac{(t-2)!}{(i-1)!(t-i-1)!} C(i,d)(t-i-1)! \\
&= 2 \sum_{i=d}^{t-1} \frac{(t-2)!}{(i-1)!} C(i,d) \\
&= 2(t-2)! \sum_{i=d}^{t-1} \frac{C(i,d)}{(i-1)!} \\
&= 2(t-2)! \sum_{i=d}^{t-1} \frac{C(i,d)}{C(i)}.
\end{aligned}$$

There is one $t$-edge per tree; therefore, $C(t,t)$ is simply $C(t)$.

#### G Read likelihood

The likelihood of the reads given the cell's genotype, base-calling error probabilities, read coverage, amplification, and allelic dropout probabilities is

$$
\begin{aligned}
&P(\mathbf{R}_c|X_c, \mathbf{Q}_c, L_c, p_{ae}, p_{ado}) \\
&\quad = \sum_{D_c^1=0}^{1} \sum_{D_c^2=0}^{1} P(D_c^1, D_c^2|p_{ado}) \sum_{A_c=0}^{1} P(A_c|D_c^1, D_c^2, L_c, p_{ae}) \\
&\qquad \sum_{F_c,N_c} P(F_c, N_c|D_c^1, D_c^2, A_c, X_c, L_c) P(\mathbf{R}_c|F_c, N_c, \mathbf{Q}_c).
\end{aligned}
$$

The ADO events are modeled as two independent Bernoulli distributions with the same success probability $p_{ado}$;

$$
\begin{aligned}
P(D_c^1, D_c^2|p_{ado}) &= P(D_c^1|p_{ado})\ P(D_c^2|p_{ado}) \\
&= Be(D_c^1|p_{ado})\ Be(D_c^2|p_{ado}) \\
&= p_{ado}^{D_c^1+D_c^2}\ (1-p_{ado})^{2-(D_c^1+D_c^2)}.
\end{aligned}
$$

The number of edges in the amplification trees depends on the ADO events and the number of observations, $L_c$. An example is illustrated in Fig 3.

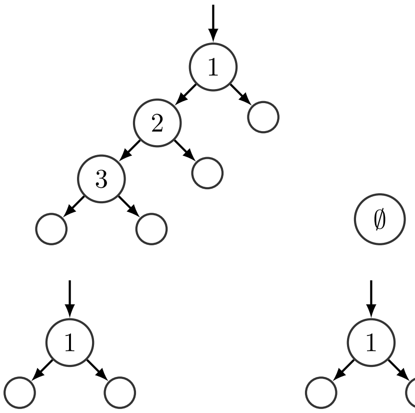


**Fig 3. Illustration of different amplification processes with $L_c = 4$ observations. Top row:** An example where the second allele is dropped ($D_c^1 = 0$ and $D_c^2 = 1$). **Bottom row:** An example with no allelic dropout event ($D_c^1 = 0$ and $D_c^2 = 0$).

$$
E_{D_c^1,D_c^2}^{L_c} = \begin{cases} 2L_c - 2, & \text{if } D_c^1 = 0, D_c^2 = 0 \\ 2L_c - 1, & \text{if } D_c^1 = 0, D_c^2 = 1 \\ 2L_c - 1, & \text{if } D_c^1 = 1, D_c^2 = 0 \\ 0, & \text{if } D_c^1 = 1, D_c^2 = 1. \end{cases}
$$

The probability of the number of AEs is a Binomial distribution of $E_{D_c^1,D_c^2}^{L_c}$ trials with $p_{ae}$ success probability;

$$
\begin{aligned}
P(A_c|D_c^1, D_c^2, L_c, p_{ae}) &= Bin(A_c|E_{D_c^1,D_c^2}^{L_c}, p_{ae}) \\
&= \binom{E_{D_c^1,D_c^2}^{L_c}}{A_c} p_{ae}^{A_c}\ (1-p_{ae})^{E_{D_c^1,D_c^2}^{L_c} - A_c}.
\end{aligned}
$$

Since the AE has a low probability, it is unlikely to observe more than one AE. Therefore, we only consider the cases where $A_c \in \{0, 1\}$ rather than $A_c \in [0, E^{L_c}_{D^1_c, D^2_c}]$. Then the probability of the AE count is

$$P(A_c|D^1_c, D^2_c, L_c p_{ae}) = \begin{cases} (1-p_{ae})^{E^{L_c}_{D^1_c, D^2_c}}, & \text{if } A_c = 0 \\ E^{L_c}_{D^1_c, D^2_c}\, p_{ae}\, (1-p_{ae})^{E^{L_c}_{D^1_c, D^2_c}-1}, & \text{if } A_c = 1 \\ 0, & \text{otherwise.} \end{cases}$$

The read probabilities given the fragment genotypes, fragment counts, and base-calling error probabilities are computed with dynamic programming, described in the Appendix.

Finally, the fragment probabilities are calculated, as shown in Tables 1 and 2. Table 1 shows possible combinations of random variables and associates them with a unique ID; Table 2 shows the fragment configuration probabilities corresponding to the configurations in Table 1. The columns $D^1_c$, $D^2_c$, and $A_c$ are the random variables the probability is conditioned on. $F_c$ and $N_c$ columns are the valid fragment genotype and fragment count contributions. For brevity, we omitted two significant, repetitive constraints in the table; (i) $(F^1_c, F^2_c) = X_c$ and (ii) $N^1_c + N^2_c + N^3_c = L_c$. All configurations that do not satisfy these conditions have 0 probability. In the case of AE events ($A_c = 1$), the fragment genotype that had the error is shown in the $pa(F^3_c)$ column. Finally, the fragment probabilities are displayed.

**Table 1.** Random variable configurations and associated case IDs of fragment configuration probabilities. For brevity, the conditions $(F^1_c, F^2_c) = X_c$ and $N^1_c + N^2_c + N^3_c = L_c$ are not shown but are assumed to be correct. The fragment configuration probability is simply zero if these conditions are not met.

| Case ID | $D^1_c$ | $D^2_c$ | $A_c$ | $F_c = (F^1_c, F^2_c, F^3_c)$ | $N_c = (N^1_c, N^2_c, N^3_c)$ | $pa(F^3_c)$ |
|---|---|---|---|---|---|---|
| 0 | 0 | 0 | 0 | $F_3 = \emptyset$ | $(>0,\ >0,\ 0)$ | - |
| 1 | 0 | 1 | 0 | $F_3 = \emptyset$ | $(L_c,\ 0,\ 0)$ | - |
| 2 | 1 | 0 | 0 | $F_3 = \emptyset$ | $(0,\ L_c,\ 0)$ | - |
| 3 | 1 | 1 | 0 | $F_3 = \emptyset$ | $(>0,\ >0,\ 0)$ | - |
| 4 | 0 | 0 | 1 | $d(F_3\|\|F_1) = 1,\ d(F_3\|\|F_2) \neq 1$ | $(\geq 0,\ >0,\ >0)$ | $F_1$ |
| 5 | 0 | 0 | 1 | $d(F_3\|\|F_1) \neq 1,\ d(F_3\|\|F_2) = 1$ | $(>0,\ \geq 0,\ >0)$ | $F_2$ |
| 6 | 0 | 0 | 1 | $d(F_3\|\|F_1) = 1,\ d(F_3\|\|F_2) = 1$ | $(0,\ >0,\ >0)$ | $F_1$ |
| 7 | 0 | 0 | 1 | $d(F_3\|\|F_1) = 1,\ d(F_3\|\|F_2) = 1$ | $(>0,\ 0,\ >0)$ | $F_2$ |
| 8 | 0 | 0 | 1 | $d(F_3\|\|F_1) = 1,\ d(F_3\|\|F_2) = 1$ | $(>0,\ >0,\ >0)$ | $F_1$ or $F_2$ |
| 9 | 0 | 1 | 1 | $d(F_3\|\|F_1) = 1$ | $(\geq 0,\ 0,\ >0)$ | $F_1$ |
| 10 | 1 | 0 | 1 | $d(F_3\|\|F_2) = 1$ | $(0,\ \geq 0,\ >0)$ | $F_2$ |

**Table 2.** Case IDs and corresponding fragment configuration probabilities. For brevity, the conditions $(F_c^1, F_c^2) = X_c$ and $N_c^1 + N_c^2 + N_c^3 = L_c$ are not shown but are assumed to be correct. The fragment configuration probability is simply zero if these conditions are not met.

| Case ID | $p(F_c, N_c \| D_c^1, D_c^2, A_c, X_c, L_c)$ |
|---|---|
| 0 | $\frac{1}{L-1}$ |
| 1 | 1 |
| 2 | 1 |
| 3 | 1 |
| 4 | $\frac{1}{L-1}\frac{C(N_1+N_3,N_3)}{C(N_1+N_3)}\frac{1}{E^L_{D_1,D_2}}\frac{1}{6}$ |
| 5 | $\frac{1}{L-1}\frac{C(N_2+N_3,N_3)}{C(N_2+N_3)}\frac{1}{E^L_{D_1,D_2}}\frac{1}{6}$ |
| 6 | $\frac{1}{L-1}\frac{C(N_1+N_3,N_3)}{C(N_1+N_3)}\frac{1}{E^L_{D_1,D_2}}\frac{1}{6}$ |
| 7 | $\frac{1}{L-1}\frac{C(N_2+N_3,N_3)}{C(N_2+N_3)}\frac{1}{E^L_{D_1,D_2}}\frac{1}{6}$ |
| 8 | $\frac{1}{L-1}\left(\frac{C(N_1+N_3,N_3)}{C(N_1+N_3)}+\frac{C(N_2+N_3,N_3)}{C(N_2+N_3)}\right)\frac{1}{E^L_{D_1,D_2}}\frac{1}{6}$ |
| 9 | $\frac{C(L,N_3)}{C(L)}\frac{1}{E^L_{D_1,D_2}}\frac{1}{6}$ |
| 10 | $\frac{C(L,N_3)}{C(L)}\frac{1}{E^L_{D_1,D_2}}\frac{1}{6}$ |

### H Differences between singleton and paired sites

Here, we compiled the differences between singleton and paired sites in various equations.

- Singleton sites consist of one base pair in the genome. Paired sites consist of a pair of base pairs, one base pair is the candidate mutation site, and the other is the gSNV locus.
- The mutation type random variable, $Z$, has $K = 3$ categories for singleton sites and $K = 12$ for paired sites. The number of categories affects Eq 4 in the main manuscript.
- The third fragment type probability, $F_c^3$, is 1/3 for singleton sites and 1/6 for paired sites. Table 2 contains this probability.
- The computation of the likelihood of a selected site depends on the site type; see Eq 5 and 6 in the main manuscript.
- For the real data processing and site selection, the data in Mpileup format is sufficient for singleton site analysis since one can obtain the nucleotides and their associated Phred quality scores. On the contrary, the analysis-ready BAM files are needed for the paired sites to extract the reads covering both loci. However, a Mpileup file can be used as a guide to speed up site selection.

### I Real data preprocessing

We followed a standard pipeline to process the raw unmapped reads (bulk and single-cell DNA sequencing data in FASTQ format). The adapters are removed from the reads using Cutadapt [1]. The reads are mapped to the GRCh37 human reference genome using Bowtie2 [2, 3]. The mapped reads are converted to BAM format, and the duplicate reads are marked using Picard [4]. The reads are realigned based on the known indels (1000 Genomes Phase I and Mills and 1000 Genomes Gold Standard Indels) using GATK [5]. The base quality scores are recalibrated using GATK, and analysis-ready reads in BAM format are obtained.

In order to identify gSNV sites, we used FreeBayes [6] software on bulk data. The minimum alternate count is set to 10, and the minimum alternate fraction is set to 0.2. The reported heterozygous SNPs (0/1) are used as gSNV sites. The regions around gSNV sites are used for analysis.

SCIΦ requires the input data to be in Mpileup format.[1] Samtools [7] is used to pile up individual BAM files.

[1] Our software uses the Mpileup format for faster site detection, which is not a mandatory file format.

#### J Fibroblast dataset information

Table 3 shows the single-cell ids, their clonal information, and approximate read coverages. For more information, see [8].

**Table 3.** Fibroblast dataset information

| Paper Cell ID | Donor | Clone ID | Project Cell ID | Coverage |
|---|---|---|---|---|
| 0 | C5RO | 1 | 22 | 15x |
| 1 | C5RO | 1 | 24 | 15x |
| 2 | C5RO | 1 | 27 | 15x |
| 3 | C5RO | 1 | 30 | 15x |
| 4 | C5RO | 1 | 33 | 15x |
| 5 | C5RO | 1 | 34 | 15x |
| 6 | C5RO | 1 | 36 | 15x |
| 7 | C5RO | 1 | 37 | 15x |
| 8 | C5RO | 1 | 38 | 15x |
| 9 | C5RO | 1 | 40 | 15x |
| 10 | C5RO | 1 | 42 | 15x |
| 11 | C5RO | 1 | 43 | 15x |
| 12 | C5RO | 2 | 4 | 10x |
| 13 | C5RO | 2 | 6 | 10x |
| 14 | C5RO | 2 | 16 | 10x |
| 15 | C5RO | 2 | 17 | 10x |
| 16 | C5RO | 2 | 19 | 10x |
| 17 | C5RO | 2 | 21 | 10x |

### K Number of sites in biological data experiments

Table 4 shows the number of sites used during the biological data experiments. The *gSNV* column is the number of bulk heterozygous sites detected by the FreeBayes [6] software. The *paired*, *singleton*, and *total* sites are the paired, singleton, and the total number of sites used by the proposed method. The SCIΦ column is the number of mutations reported by SCIΦ.

**Table 4.** The number of sites per chromosome used during the biological data experiment

| Chr | gSNV | paired | singleton | total | SCIΦ |
|---|---|---|---|---|---|
| chr 1 | 158304 | 21040 | 278844 | 299884 | 213796 |
| chr 2 | 168536 | 18995 | 264076 | 283071 | 188241 |
| chr 3 | 148956 | 16957 | 222185 | 239142 | 152300 |
| chr 4 | 145413 | 13781 | 196315 | 210096 | 127486 |
| chr 5 | 134734 | 12274 | 190120 | 202394 | 134922 |
| chr 6 | 131005 | 15957 | 214361 | 230318 | 150313 |
| chr 7 | 116606 | 12754 | 175727 | 188481 | 123977 |
| chr 8 | 106457 | 12970 | 174987 | 187957 | 121738 |
| chr 9 | 86655 | 9916 | 132501 | 142417 | 97515 |
| chr 10 | 104292 | 12634 | 168017 | 180651 | 120637 |
| chr 11 | 106351 | 12233 | 171426 | 183659 | 122519 |
| chr 12 | 99553 | 10896 | 151332 | 162228 | 101703 |
| chr 13 | 75318 | 5854 | 91960 | 97814 | 64348 |
| chr 14 | 65728 | 6956 | 100771 | 107727 | 73194 |
| chr 15 | 58000 | 7559 | 96954 | 104513 | 74673 |
| chr 16 | 69048 | 10161 | 113141 | 123302 | 84303 |
| chr 17 | 59305 | 8673 | 103650 | 112323 | 76798 |
| chr 18 | 56471 | 6363 | 79423 | 85786 | 63121 |
| chr 19 | 45606 | 6026 | 77229 | 83255 | 61581 |
| chr 20 | 46514 | 8997 | 93367 | 102364 | 70059 |
| chr 21 | 31124 | 4700 | 44346 | 49046 | 29857 |
| chr 22 | 27352 | 5721 | 61414 | 67135 | 50133 |
| Total | 2041328 | 241417 | 3202146 | 3443563 | 2303214 |

#### L Similarity score comparison of all methods

In this section, we compare the similarity score of all four methods, Scuphr, Scuphr with default parameters ($p_{ado} = 0.1$ and $p_{ae} = 0.01$), SCIΦ, and SCIΦ with candidate sites selected by Scuphr. Fig 4 shows the similarity scores of all methods in the low AE dataset, and Fig 5 shows the similarity scores of all methods in the high AE dataset.

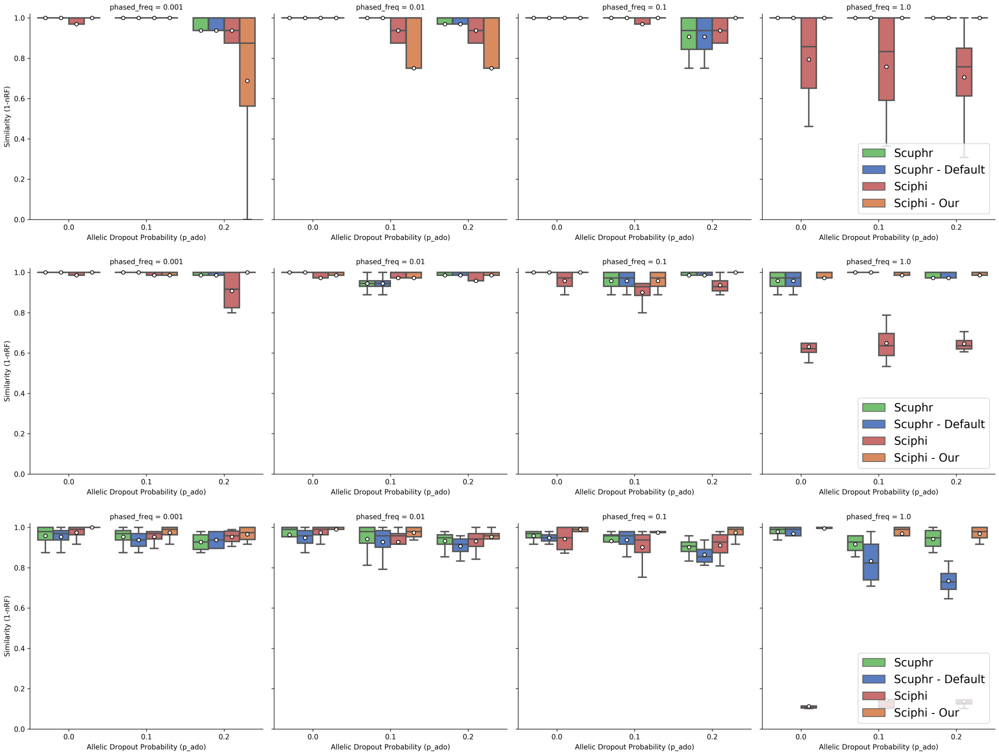


**Fig 4. Similarity scores of all methods for low amplification error datasets. Top row:** Results for 10 cells. **Center row:** Results for 20 cells. **Bottom row:** Results for 50 cells.

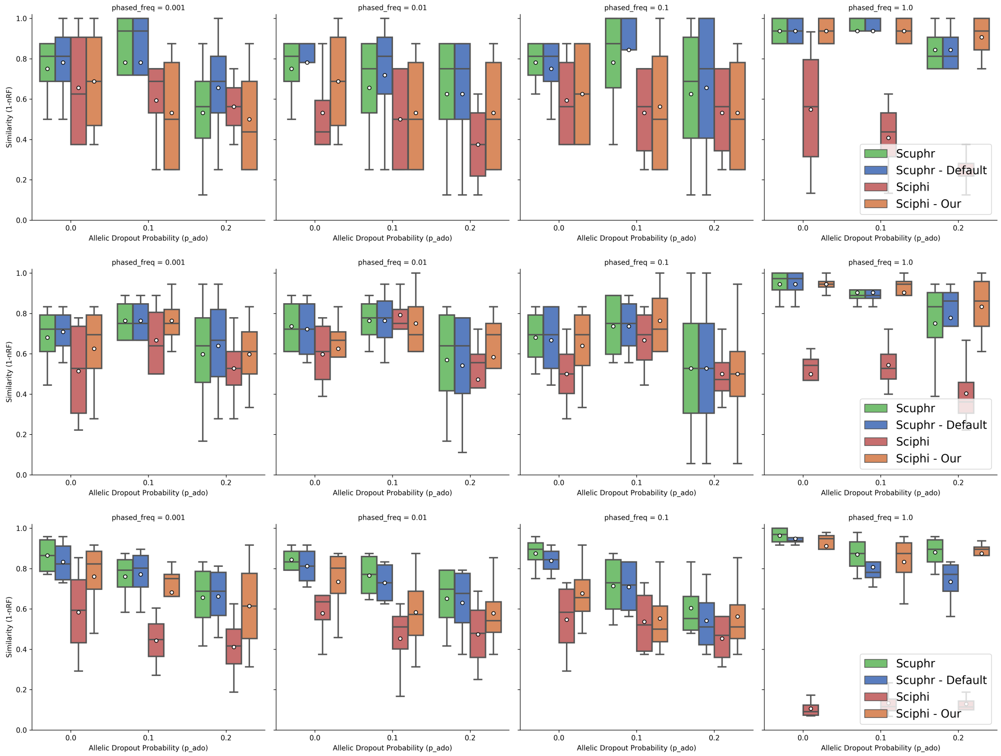


**Fig 5. Similarity scores of all methods for high amplification error datasets.** **Top row:** Results for 10 cells. **Center row:** Results for 20 cells. **Bottom row:** Results for 50 cells.

### M Runtime analysis for parameter estimation

Parameter estimations are done using 20 singleton and paired sites. Each parameter estimation is performed by running three independent chains for $5,000$ iterations sequentially. Each configuration is repeated ten times, and the results are shown in Fig 6 and 7. The runtimes increase with the number of cells. Parameter estimation with the paired sites requires more wall-clock time than singleton sites. Scuphr saves intermediate states and reuses them frequently instead of recomputing the same states. The parallelization is done per site; hence no substantial performance gains are going from 16 to 32 cores.

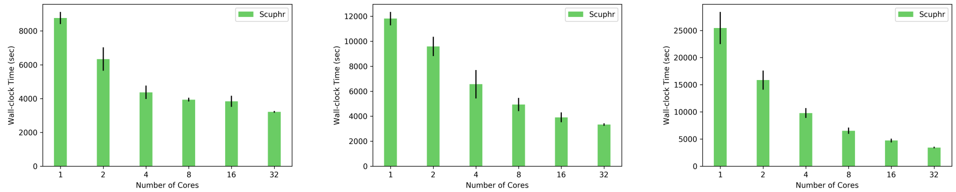


**Fig 6. Runtime comparison of parameter estimation using singleton sites.** The x-axis is the number of cores, and the y-axis is the wall-clock time in seconds. Standard deviations are shown with vertical lines. **Left:** Runtime for 10 cells. **Center:** Runtime for 20 cells. **Right:** Runtime for 50 cells.

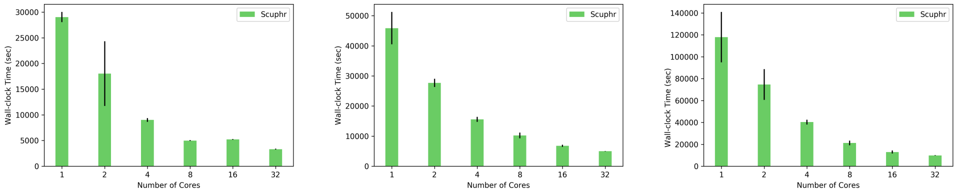


**Fig 7. Runtime comparison of parameter estimation using paired sites.** The x-axis is the number of cores, and the y-axis is the wall-clock time in seconds. Standard deviations are shown with vertical lines. **Left:** Runtime for 10 cells. **Center:** Runtime for 20 cells. **Right:** Runtime for 50 cells.

### N Runtime comparison with more cores for SCIΦ

In addition to the single-core SCIΦ runs, we compared the runtimes with multiple-core SCIΦ runs for a subset of experiment configurations, as presented in Fig 8. As a result of the embarrassingly parallel distance matrix computation, our method scales linearly with the number of cores. Even though SCIΦ is not an inherently parallel method, we observed slight performance gains with the increasing number of cores. However, this gain is not linear, and we hypothesize that the gain is primarily due to the efficient computations of used libraries in the software.

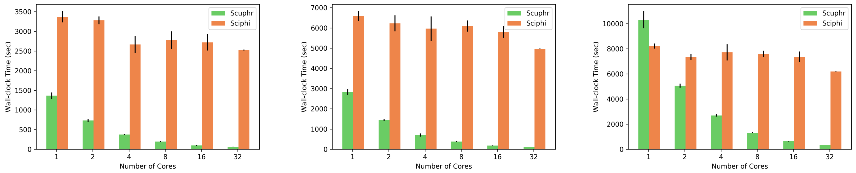


**Fig 8. Runtime comparison of SCIΦ with the varying number of cores for singleton sites.** The x-axis is the number of cores, and the y-axis is the wall-clock time in seconds. Standard deviations are shown with vertical lines. Left, center, and right subplots are the results for the (cell, site) tuples (20, 256), (20, 512), and (50, 256), respectively.

#### O SCIΦ details

We installed SCIΦ version v0.1.7 using Bioconda [9]. In all of the experiments, the software's default parameters are used. The bulk data is provided as *control bulk normal* (BN), and the single-cell data are inputted as *tumor cells* (CT). The MCMC chains are run for 1,100,000 iterations, and the reported tree is used for analysis.

For the synthetic datasets with high phasing frequency (e.g., 1), SCIΦ picked a small number of sites for analysis due to high heterogeneity in the genome. We did additional experiments and provided the sites our software picked using the SCIΦ software's inclusion and exclusion lists features. We used Monovar [10] to detect variants and picked the reference and alternate nucleotide information of inclusion sites from its output. For the sites Monovar did not detect, we picked the most common non-reference nucleotide as the alternate.

For the real dataset, SCIΦ is applied to each chromosome independently (due to the file sizes of Mpileup format). The resulting trees are sampled with replacement, weighted with respect to the number of mutations, for bootstrapping.
